# Decoding the IGF1 Signaling Gene Regulatory Network Behind Alveologenesis from A Mouse Model of Bronchopulmonary Dysplasia

**DOI:** 10.1101/2022.01.24.477613

**Authors:** F Gao, C Li, SM Smith, N Peinado, G Kohbodi, E Tran, E Loh, W Li, Z Borok, P Minoo

## Abstract

Lung development is precisely controlled by underlying Gene Regulatory Networks (GRN). Disruption of genes in the network can interrupt normal development and cause diseases such as bronchopulmonary dysplasia (BPD)–a chronic lung disease in preterm infants with morbid and sometimes lethal consequences characterized by lung immaturity and reduced alveolarization.

Here, we generated a transgenic mouse exhibiting a moderate severity BPD phenotype by blocking IGF1 signaling in secondary crest myofibroblasts (SCMF) at the onset of alveologenesis. Using approaches mirroring the construction of the model GRN in sea urchin’s development, we constructed the IGF1 signaling network underlying alveologenesis using this mouse model that phenocopies BPD. The constructed GRN, consisting of 43 genes, provides a bird’s-eye view of how the genes downstream of IGF1 are regulatorily connected. The GRN also reveals a mechanistic interpretation of how the effects of IGF1 signaling are transduced within SCMF from its specification genes to its effector genes and then from SCMF to its neighboring alveolar epithelial cells with WNT5A and FGF10 signaling as the bridge. Consistently, blocking WNT5A signaling in mice phenocopies BPD as inferred by the network. A comparative study on human samples suggests that a GRN of similar components and wiring underlies human BPD.

Our network view of alveologenesis is transforming our perspective to understand and treat BPD. This new perspective calls for the construction of the full signaling GRN underlying alveologenesis, upon which targeted therapies for this neonatal chronic lung disease can be viably developed.

## Introduction

Development is precisely controlled by the genetic program encoded in the genome and is governed by genetic interactions (Davidson et al., 2002; Levine and Davidson, 2005). Elucidating the network of interactions among genes that govern morphogenesis through development is one of the core challenges in contemporary functional genomics research (Przybyla and Gilbert, 2021). These networks are known as developmental gene regulatory networks (GRN) and are key to understanding the developmental processes with integrative details and mechanistic perspectives. Over the past few decades, great advances have been made in decoding these networks in classical model systems (i.e. Dequeant and Pourquie, 2008; Longabaugh et al., 2017; Olson, 2006; Satou et al., 2009; Sauka-Spengler and Bronner-Fraser, 2008), as well as in forming in-depth understandings of the general design principles of these networks in development and evolution (Carre et al., 2017; Davidson, 2010; Erwin and Davidson, 2009; Gao and Davidson, 2008; Lim et al., 2013; Peter and Davidson, 2009; Royo et al., 2011). Highlighted among them is the sea urchin developmental GRN, the most comprehensive and authenticated network constructed to date (Peter and Davidson, 2017).

Disruption of any gene in the network could interrupt normal development and conceivably cause diseases. Some diseases are caused by single gene disorders while others are complex and multifactorial such as bronchopulmonary dysplasia (BPD), a common cause of morbidity and mortality in preterm infants characterized by arrested alveolar development with varied severities within the subjects’ lungs (Jain and Bancalari, 2014; Jobe, 1999; Short et al., 2007). Both prenatal insults and postnatal injury increase the risk of BPD. The multifactorial etiology of BPD has made the development of therapies a unique challenge, and currently, no effective treatment exists to prevent or cure this debilitating disease.

In recent clinical trials, therapy with recombinant human insulin-like growth factor 1 (rhIGF1) protein showed initial promise for BPD, but significant limitations did not allow further clinical trials (Ley et al., 2019; Seedorf et al., 2020). One impediment to further development of IGF1 as a therapy is the lack of comprehensive information regarding the precise role of the IGF1 signaling pathway in alveologenesis.

Alveolar development is a highly complex process that is driven by multiple signaling pathways including TGFB, SHH, WNT, PDGF, VEGF, HGF, NOTCH, BMP, and IGF1 (Juul et al., 2020; Nabhan et al., 2018; Tsao et al., 2016; Verheyden and Sun, 2020; Wu and Tang, 2021; Zepp and Morrisey, 2019). IGF1 and IGF1R expression is found throughout fetal lung development and fluctuates at different stages. Both *IGF1* and *IGF1R* expression is significantly reduced in BPD lungs (Hellstrom et al., 2016; Lofqvist et al., 2012; Yilmaz et al., 2017), suggesting a potential role in the pathogenesis of BPD. The function of IGF1 and IGF1R has been examined in their respective constitutive and conditional knockout mice, but their specific function during alveologenesis has not been examined (Epaud et al., 2012; Lopez et al., 2016; Lopez et al., 2015). A recent study found interruption in IGF1 signaling compromised mechanosignaling and interrupted alveologenesis (He et al., 2021).

In our study here, we found *Igf1* and *Igf1r* are primarily expressed in SCMF in postnatal mice lungs. As a result, we interrupted IGF1 signaling in SCMF by the inactivation of *Igf1r* at the onset of alveologenesis in postnatal day 2 (PN2) mouse neonates and analyzed the resulting phenotypes. Inactivation of *Igf1r* resulted in mutant lungs with simplified and immature alveolar structure resembling that of human BPD at a moderate severity (Short et al., 2007). We reasoned that the phenotype caused by the inactivation of *Igf1r* in SCMF reflects interruption in the genetic program downstream of IGF1 signaling that drives the normal functions of SCMF in the process of alveologenesis. Alterations in SCMF may further impact specification/differentiation of other key cell types, particularly the alveolar epithelial cells. We aim to decode the genes and their interactions behind this underlying genetic program.

Taking advantage of the high throughput next generation sequencing, people have been trying to build GRNs by computational analysis using wild type gene expression data (i.e. Jia et al., 2017; Xu et al., 2012). Currently, the most reliable way is still to build GRN experimentally on data from perturbation analyses.

Genes we particularly focus on are the regulatory genes. These genes decide the outcome of a GRN as they regulate and control other genes’ expression forming a regulatory circuitry of varied hierarchy with structural and cellular genes as their terminal targets (Davidson, 2010; Erwin and Davidson, 2009). Their functions are defined by their logical control over the circuitry’s operation and their synergetic biological effects are manifested by the effector genes under the circuitry (Peter and Davidson, 2009; Peter et al., 2012).

We followed the protocol similar to that used in the construction of the sea urchin GRN (Materna and Oliveri, 2008): regulatory genes were selected at the transcriptomic scale from the LungMAP database (www.lungmap.net) where their expression during alveologenesis was reported; the cellular expression patterns of these genes were then annotated on lungMAP scRNAseq data for the screen of SCMF genes; the regulatory interactions among these genes were subsequently examined from in vivo and in vitro perturbations; and cellular communications were finally determined by Secretome-Receptome analysis. Combining this data, we constructed the IGF1 signaling gene regulatory network underlying alveologenesis from this mouse model of human BPD phenocopy.

Our GRN work on alveologenesis represents a transformative view of how to understand and perhaps even design future preventive and therapeutic strategies for BPD treatment.

## Results

### Postnatal Expression of *Igf1* and *Igf1r* in the lung

To define the expression pattern of *Igf1* and *Igf1r* during early postnatal lung development, we performed RT-PCR using total lung RNA from embryonic day E18 to postnatal day PN30. The analysis showed *Igf1* is expressed dynamically during development, peaking at the onset of alveologenesis and progressively decreasing thereafter (Fig.1A). The expression pattern of *Igf1r* was biphasic, initially displaying an overlap with *Igf1* in early alveologenesis with a subsequent peak occurring on PN21 (Fig.1B). Expression profiling of Igf1 and Igf1r from the entire lung and within Myofibroblasts during lung development was calculated based on data from the LungMAP (Figure 1-Figure Supplement 1A) and from the latest scRNAseq dataset (Figure 1-Figure Supplement 1B, Negretti et al., 2021). Similar patterns of their expression were observed when compared to our data.

**Figure 1:**
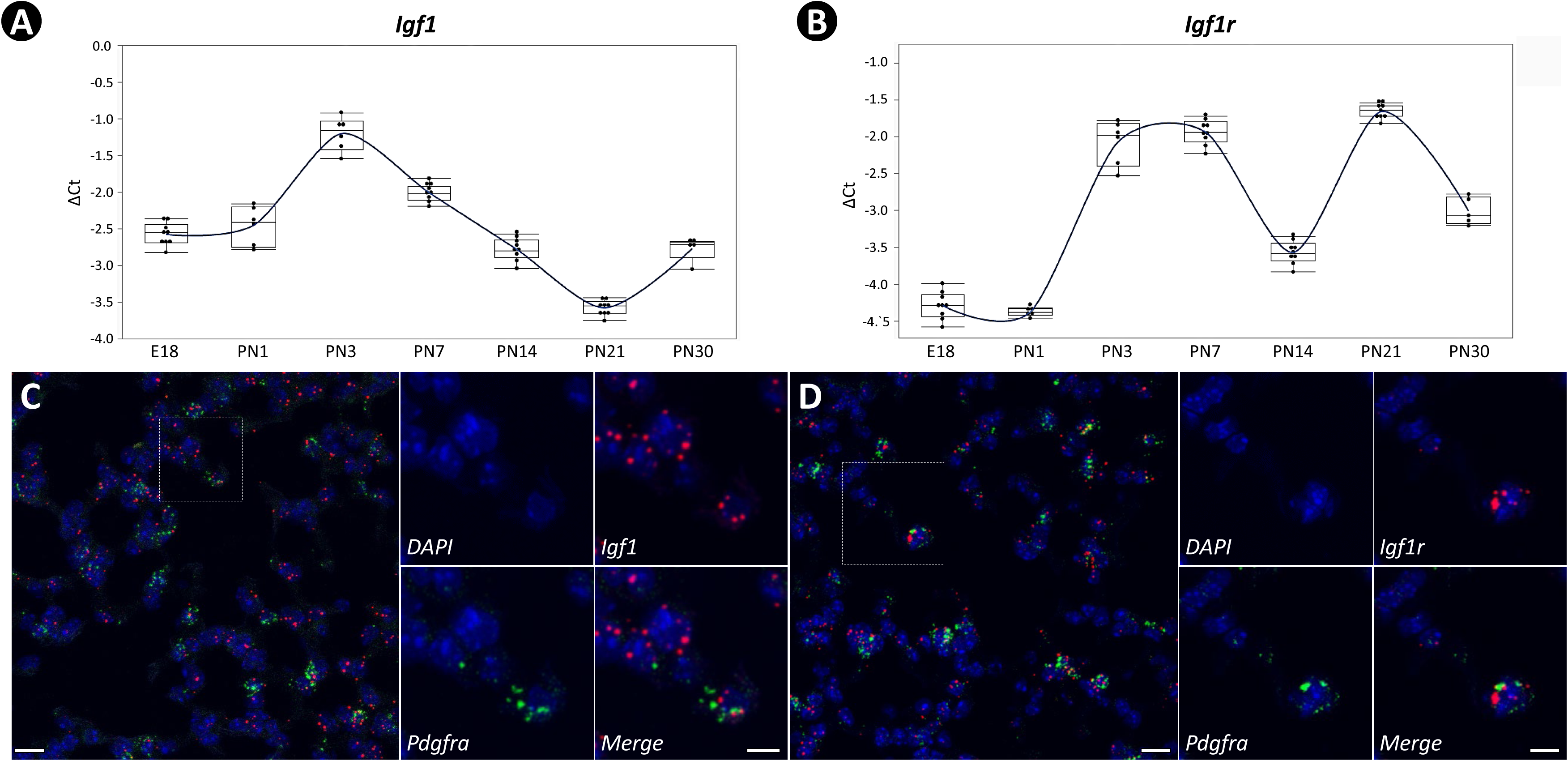
Temporal and spatial expression of *Igf1* and *Igf1r* during neonatal lung development in mice. A, B: Temporal expression of *Igf1* (A) and *Igf1r* (B) from E18 to PN30, quantified by RT-PCR and normalized to *Gapdh*. RNA used was collected from the whole lung and each stage was represented by at least five lungs. C: Spatial localization of mRNA for *Igf1* (C) and *Igf1r* (D) in PN7 lungs as detected by RNAscope and their overlapped expression with *Pdgfra*. Outlined area on the left is magnified on the right. Scale bars: 20um under the whole view and 10um under the magnified view.

Alveologenesis is characterized by secondary septa/crest formation, and the Alveolar Myofibroblasts are recognized as the driving force behind it. Secondary Crest Myofibroblast was derived from this concept and is broadly adopted in independent publications (i.e. Bostrom et al., 1996; Li et al., 2015; Li et al., 2018; Sun et al., 2022; Zepp et al., 2021).

The definition, specification, and function of SCMFs have not been systematically characterized. Presently, a consensus marker for these cells hasn’t been established although a list of markers, such as *Acta2* (Kugler et al., 2017), *Stc1* (Zepp et al., 2021), *Tagln* (Li et al., 2018), *Fgf18* (Hagan et al., 2020) and *Pdgfra* (Li et al., 2015) have been suggested from previous work. To help resolve this issue, we compiled a comprehensive list of these markers suggested from literature and examined their expression across different mesenchymal cell types as clustered on the two latest scRNAseq datasets (Figure 1-Figure Supplement 1C&F, Negretti et al., 2021, Zepp et al., 2021). *Pdgfra* was revealed as a good SCMF marker agreed by both datasets (Figure 1-Figure Supplement 1C&F). RNAscope showed highly overlapping localization of *Igf1/Igf1r* and *Pdgfra* (Fig.1C&D), indicating SCMF as a principal cellular site of *Igf1* and *Igf1r* expression in lungs undergoing alveologenesis.

### Mesodermal-specific Inactivation of *Igf1* and *Igf1r*

As SCMF are derived from the lung mesenchyme early in lung development and *Dermo1* is activated at the onset of mesodermal lineage specification (Li et al., 2008), we used *Dermo1-cre* (*Dcre*) to inactivate *Igf1* and *Igf1r* separately, specifically within mesodermal lineages, and examined the mutant lung phenotype during embryonic and postnatal development (Figure 2-Figure Supplement 1A).

Loss of *Igf1* or *Igf1r* in mesodermal progenitors using floxed alleles of these genes decreased their mRNA by approximately two folds (Figure 2-Figure Supplement 1B). Consistent with previous reports (Epaud et al., 2012; Lopez et al., 2015), reduction in body weight was observed in our *Igf1r* mutant pups (Figure 2-Figure Supplement 1C&D).

The *Dcre;Igf1r-f/f* mutant displayed clearly visible lung defects: thickened saccular walls at E18; dilated sacculi and severe reductions in the number of secondary crests at PN14 with the latter extending into PN30 (Figure 2-Figure Supplement 1E). Similar defects of less severity were observed in the *Dcre;Igf1-f/f* mutant (Figure 2-Figure Supplement 1F). However, due to the timing of *Dermo1-cre* activation, it is difficult to determine whether the impact of *Igf1r* on alveologenesis occurs postnatally or is a carryover impact of events occurring prior to onset of alveologenesis.

### Postnatal inactivation of *Igf1r* in SCMF profoundly arrests alveologenesis

The SHH targeted (i.e. Gli1+) fibroblasts have been rigorously examined through lineage tracing using *Gli1creERT2;mTmG* in our lab (i.e. Li et al., 2015, Li et al., 2019). Cell lineage analysis shows the SHH signaling targets different mesenchymal cell lineages through lung development.

There was a window of time in the early postnatal stage during which the derived GFP+ cells were observed primarily localized to the secondary septa and secondarily to parabronchial & perivascular smooth muscle fibers (Li et al., 2015). When Tamoxifen was titrated down to a certain dosage, the smooth muscle fibers were no longer labeled by GFP (Figure 2-Figure Supplement 2C). This very specific regimen was employed in our current paper. Consistent with our observation, it was found that Gli1 is predominantly expressed by proliferative SCMF/Myofibroblast cells as calculated from the two largest scRNAseq datasets recently published (Figure 1-Figure Supplement 1D&G; Negretti et al., 2021; Zepp et al., 2021).

Using *Gli1-creERT2 (Gcre)*, we inactivated floxed alleles of *Igf1r* on PN2 at the onset of alveologenesis (Fig.2A). Inactivation of *Igf1r* was validated by genotyping (Figure 2-Figure Supplement 2A) and its downregulation verified by RT-PCR with RNA from both entire lung and FACS-isolated SCMF (Figure 2-Figure Supplement 2B). A successful recombination was assessed by Cre-induced green fluorescent protein (GFP) (Figure 2-Figure Supplement 2C). The mutant mice were slightly runted compared to the controls (Figure 2-Figure Supplement 2D).

**Figure 2:**
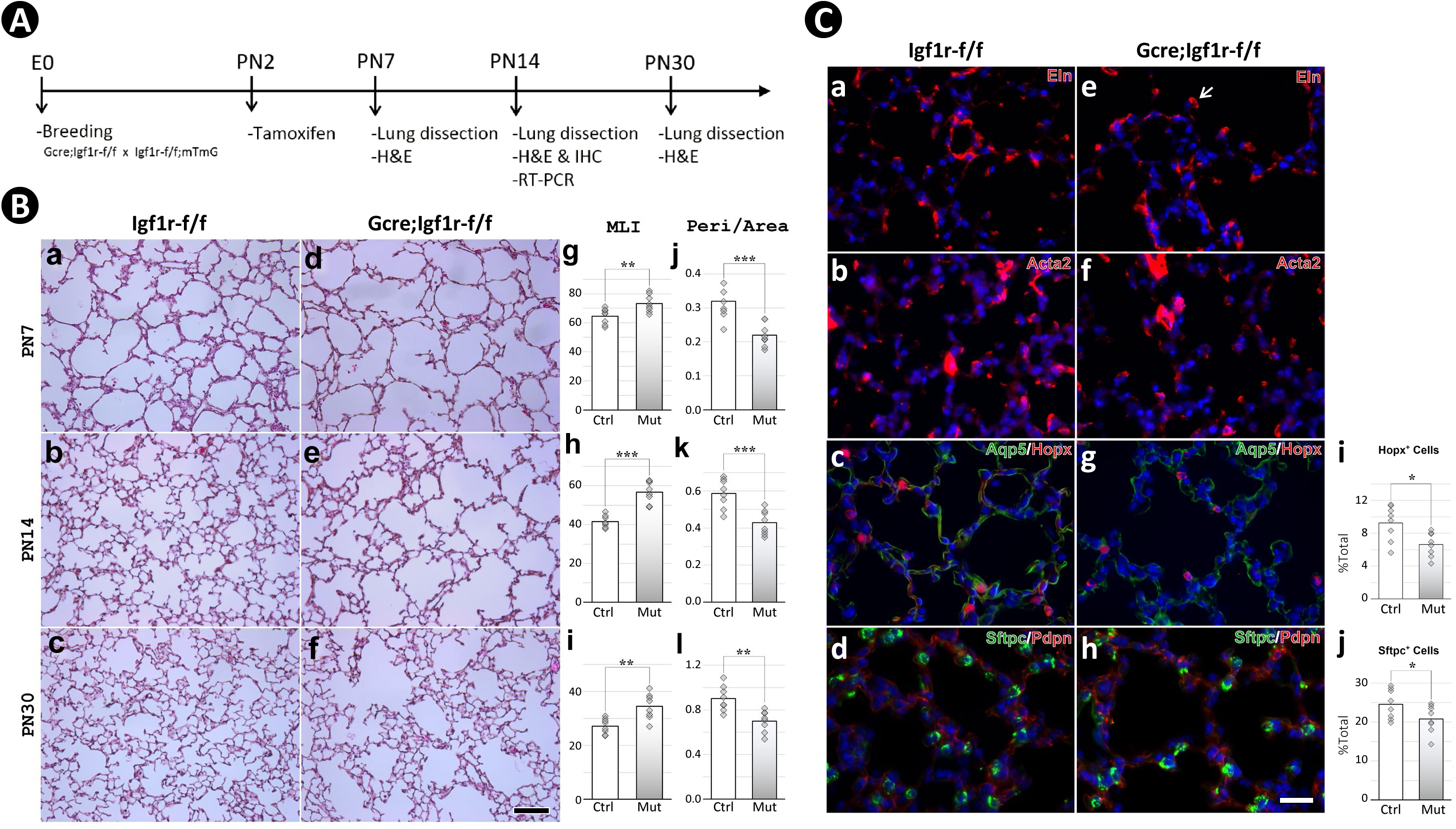
Postnatal inactivation of *Igf1r* from lung secondary crest myofibroblast cells. A: Schematic of the experimental protocol. B: H&E staining of lung sections from control (a-c) and *Gli1-cre^ERT2^* mutant (d-f) mice and their morphometric measurements by MLI (g-i) and Peri/Area ratio (j-l) at PN7, PN14 and PN30. See Figure 2-Figure Supplement 1 for the definition and calculation of these indices. C: Immunostaining of lung sections from control (a-d) and mutant (e-h) mice for Elastin (a,e), ACTA2 (b,f), AQP5/HOPX (c,g) and SFTPC/PDPN (d,h), and the comparison of the number of AT1 (i) and AT2 (j) cells between the control and mutant. Scale bar: 100um in B and 25um in C. *p*-value: * stands for 0.05-0.01, ** for 0.01-0.001, *** for <0.001, n.s. for not significant.The same designation is used throughout the paper.

Histology of multiple *Gcre;Igf1r-f/f* lungs at the timepoints PN7, PN14, and PN30, revealed a phenotype of profoundly arrested alveolar formation as measured by the mean linear intercept (MLI) (Fig.2B). In addition, ImageJ analysis showed decreased perimeter to area ratio of airspace (Peri/Area) as well as the number of airspaces per unit of area (# of Airspace/Area) – all consistent with the alveolar hypoplasia phenotype in the mutant lungs (Fig.2B, Figure 2-Figure Supplement 2E,F). The largest deviation between the controls and mutants in these measurements occurred at PN14, which marks the midpoint in the alveologenesis phase. Still, the severity of hypoplasia here is eclipsed by that seen in *Gcre;Tgfbr-f/f* induced BPD phenocopies (Gao et al., 2022).

Immunohistochemical analysis revealed no significant gross changes between the control and mutant lungs in either proliferation or apoptosis (Figure 2-Figure Supplement 2G). Similarly, examination of specific markers for various lung alveolar cell lineages, including the mesenchyme (ELN, TPM1), myofibroblasts (ACTA2, PDGFRA), endothelial cells (EMCN), lipofibroblasts (ADRP), epithelial cells (NKX2.1, SFTPC, HOPX), and pericytes (NG2) did not reveal significant differences (Fig.2C, Figure 2-Figure Supplement 2H). Nonetheless, it was observed in the mutant lungs that the deposition of elastin seemed aggregated at the secondary crest tips and the number of AT1 and AT2 cells were reduced (Fig.2C). The altered elastin deposition has been reported in previous studies (He et al., 2021; Li et al., 2019), but whether this alteration is the cause or the consequence of the impaired alveolar formation remains unknown.

To examine the genetic changes in *Gcre;Igf1r-f/f* mutant lungs, we tested two selected groups of genes: lung signature (Figure 2-Figure Supplement 2I) and angiogenesis (Figure 2-Figure Supplement 2J) genes. The analysis showed significant alterations in a few genes including *Fgf10*, *Igf1r*, and *Pdgfra* in the mutant lungs. It is noteworthy that the RNA used in the test was from whole lung tissue, while inactivation of *Igf1r* by *Gcre* was only targeted to SCMF. To determine the cell specific impacts of *Igf1r* inactivation, we characterized gene expression in FACS sorted cells in the following studies.

### Identification of SCMF genes altered in mouse lungs of BPD phenotype

Both GFP+ (SCMF) and Tomato+ (non-SCMF) cells were isolated by FACS from lungs dissected from *Gcre;mTmG* (control) and *Gcre;mTmG;Igf1r-f/f* (mutant) mice. Using the sorted cells, we compared the gene expression between them to identify genes enriched within SCMFs, those altered within SCMFs, and those altered within non-SCMFs in the mutant lungs (Fig.3A, Figure 3-Figure Supplement 1A).

**Figure 3:**
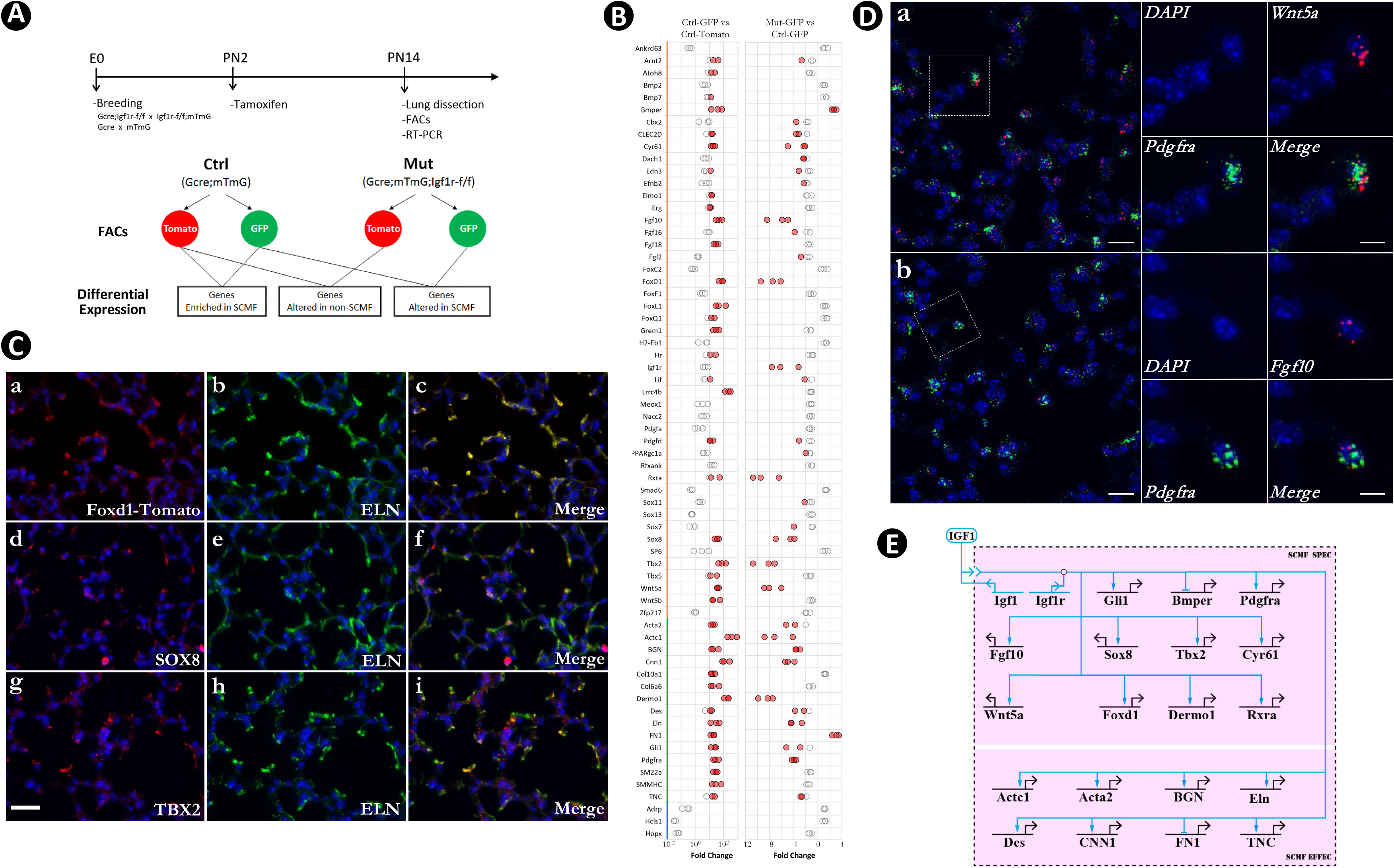
Identification of SCMF genes altered in *Gli1-cre^ERT2^;Igf1r-f/f* mutant lungs. A: Schematic of the experimental protocol. B: RT-PCR data from the selected genes displaying their enrichment in SCMF and alterations in the mutant. Genes marked with the orange line: regulatory genes selected from LungMap database; Genes with the green line: common SCMF markers; Genes with the blue line: non-SCMF genes. Red circles: data points meeting the cutoff criteria described in the text; Empty circles: data points failing the cutoff criteria. This designation is used throughout the manuscript. C: Spatial expression of Foxd1-Tomato/ELN (a-c), SOX8/ELN (d-f), and TBX2/ELN (g-i) in the alveolar compartment of PN14 lungs as detected by immunostaining. Specific antibodies were used for SOX8, TBX2, and ELN. RFP antibody was used for Foxd1-Tomato on lungs dissected from *Foxd1GCE;CAG^Tomato^* mice. Scale bar: 25um for all images. D: Spatial expression of *Wnt5a/Pdgfra* (a) and *Fgf10/Pdgfra* (b) in the alveolar compartment of PN14 lungs as detected by RNAscope. Outlined area on the left is magnified on the right. Scale bars: 20um under the whole view and 10um under the magnified view. E: Biotapestry network illustration of altered SCMF genes and their connections to IGF1 signaling within SCMF. The source of IGF1 can be both autocrine and paracrine. Genes (nodes) are shown in the territories (colored boxes) in which they are expressed. Edges show regulation by the originating upstream factor and are either positive (arrow) or repressive (bar). Signalings across cell membranes are indicated as double arrow heads.

Our analysis didn’t focus on the components of the pathway itself which are self-conserved and usually tissue-independent (Pires-daSilva and Sommer, 2003), but was instead directed at identifying genes downstream of the pathway, especially the regulatory genes through which the developmental GRN is specified (Erwin and Davidson, 2009). 47 genes known to encode signaling molecules and transcription factors were screened from the LungMAP transcriptomic database (www.lungmap.net) where they were indicated to be expressed from SCMF between PN3 to PN14 in the mouse developmental window (Figure 3-Source Data 1). Expression of the selected genes, together with 15 known SCMF cell markers and 3 non-SCMF genes (negative control), were examined and compared by quantitative RT-PCR using RNA from the sorted cells.

Three criteria were applied to identify IGF1 signaling targets within SCMF: (1) Functionally active in SCMF with its deltaCT <= 9 (relative to *Gapdh* in Ctrl_GFP, Figure 3-Source Data 1), equivalent to >= 10 copies of transcripts per cell (copies of *Gapdh* transcripts per cell based on Barber et al., 2005); (2) highly enriched in SCMF with FC >= 10, p<=0.05 (Ctrl_GFP vs Ctrl_Tomato, Fig.3B left column, Figure 3-Source Data 1); (3) significantly altered with FC >=2, p<=0.05 (Mut_GFP vs Ctrl_GFP, Fig.3B right column, Figure 3-Source Data 1).

Nineteen genes were identified that met all 3 criteria (genes highlighted red in the first column of Figure 3-Source Data 1). Spatial localization of a select number of these genes – including *Foxd1, Sox8, Tbx2, Bmper, Actc1, Wnt5a,* and *Fgf10* – in SCMF was validated by IHC (Fig.3C, Figure 3-Figure Supplement 1B) or RNAscope (Fig.3D). Consistently, *Wnt5a* was recently reported as a signature gene of SCMF (Negretti et al., 2021), and data analysis on the two latest scRNAseq datasets revealed *Wnt5a* is dominantly expressed from SCMF/Myofibroblast along with smooth muscle cells (Figure 1-Figure Supplement 1E&H).

Regulation of the latter 19 genes by Igf1 signaling was illustrated on Biotapestry (Longabaugh, 2012) as displayed in Fig.3E. Notedly, all connections were drawn directly from the signaling, whereas the regulation may happen indirectly through the cross regulation among these genes (to be examined below).

The constructed link map points to IGF1 signaling as a positive regulator for 17 out of the 19 putative downstream genes (Fig.3E). Based on their molecular and cellular functions, the 19 genes can be stratified into two separate groups. One group is comprised of regulatory genes encoding transcription factors (*Gli1, Foxd1, Tbx2, Sox8, Dermo1, Rxra*) and signaling molecules (*Fgf10, Wnt5a, Bmper, Cyr61, Pdgfra*). The second group is comprised of *Acta2, Actc1, BGN, Des, Eln, Cnn1, FN1,* and *TNC*, representing genes that encode structural & cellular molecules. Based on the GRN’s hierarchical design (Erwin and Davidson, 2009), it is most likely that IGF1 signaling first targets the regulatory genes, referred to as SCMF specifiers, whose expression subsequently targets the downstream structural and cellular genes in SCMF (SCMF effectors).

### Cross regulation of altered SCMF genes in the mouse BPD phenocopy model

Within the inventory of altered SCMF genes, as defined above, are transcription factor *Foxd1,* one of the most reduced regulatory genes from our RT-PCR measurements, and two growth factor signaling molecules, *Fgf10* and *Wnt5a,* both known to function in postnatal lung development (Chao et al., 2016; Li et al., 2020; Zhang et al., 2020). Secretome-Receptome computation using the Fantom algorithm (Ramilowski et al., 2015) revealed the cognate receptors, FGFR1/3/4 and ROR1/2, involve in FGF10 and WNT5a signaling transduction within SCMF (Fig.4B). The results were validated on bulk RNAseq (Figure 4-Figure Supplement 1A) and the latest scRNAseq datasets (Figure 4-Figure Supplement 1B,C; Figure 4-Source Data 1). WNT5A is a highly evolutionary conserved non-canonical Wnt ligand. In spite of being identified as a non-canonical ligand, it can, under certain circumstances, signal through canonical Wnt signaling pathways directly or indirectly (i.e. Mikels and Nusse, 2006). Current analysis is focused on the signaling’s non-canonical aspect and does not exclude other possibilities.

**Figure 4:**
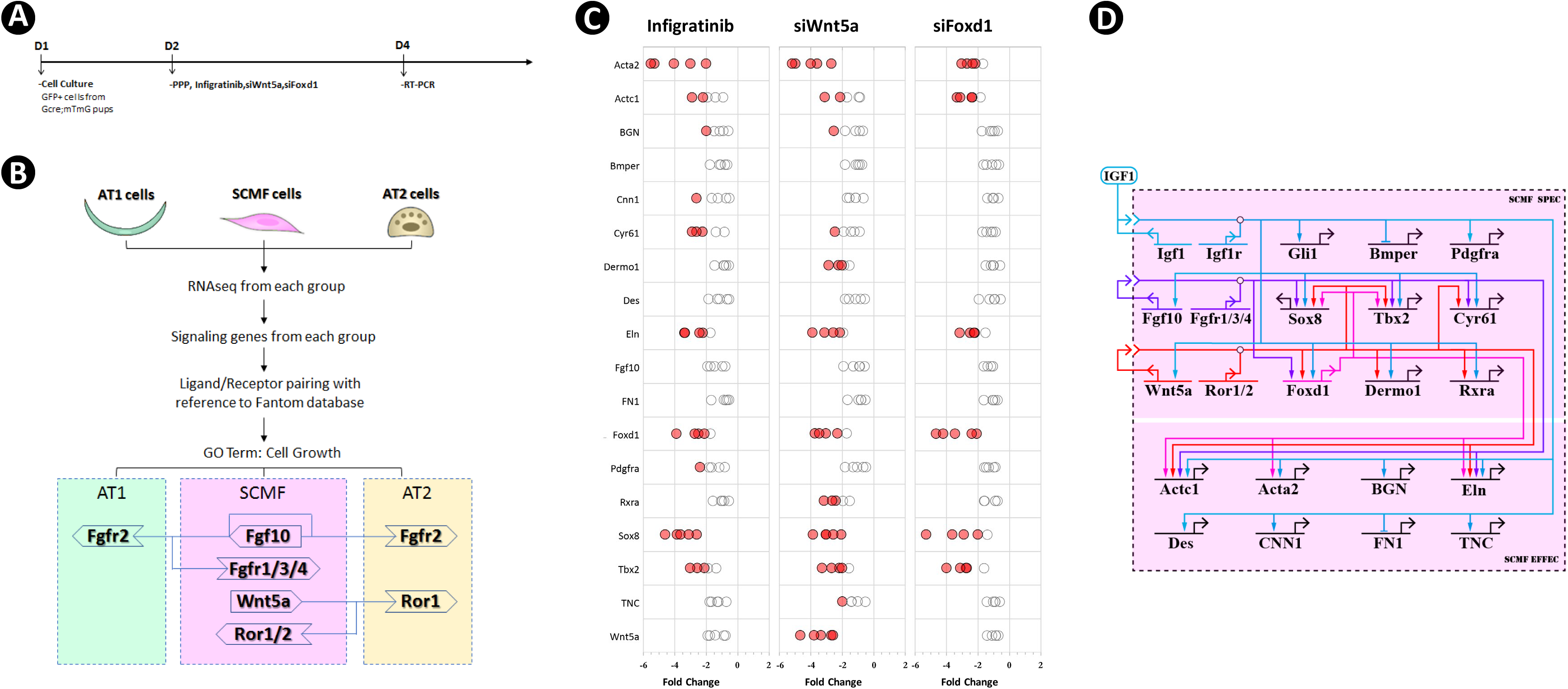
Cross regulation of the altered SCMF genes. A: Schematic of the experimental protocol. B: Flow chart of the secretome-receptome analysis among SCMF, AT1 and AT2 cells. The ligands and receptors were identified from the following RNAseq datasets: GSE126457 for SCMF (Li et al., 2019), GSE182886 for AT2, GSE106960 for AT1 (Wang et al., 2018a). C: RT-PCR data from the altered SCMF genes demonstrating their response to the treatments by Infigratinib, *siWnt5a*, and *siFoxd1*. D: Biotapestry network illustration of the cross regulation among the altered SCMF genes.

To investigate the possibility of cross regulation of these genes on the other altered SCMF genes, GFP^+^ SCMFs were isolated by FACS from postnatal *Gcre;mTmG* lungs and cultured in vitro (Fig.4 A). Cultures were treated with inhibitors which target the gene or pathway of interest. The inhibitor’s dose was determined from the literature and a series of testing (Figure 4-Source Data 2), and inhibition was validated by examining its target genes (Figure 4-Figure Supplement 2A). Blocking IGF1 signaling using the IGF1R inhibitor (PPP at 4uM as in Chen et al., 2017; Jin et al., 2018; Wang et al., 2019), led to altered expression of all genes – except *Gli1 –* in the same direction as observed in vivo (Figure 4-Figure Supplement 2B), indicating that isolated cells in vitro recapitulate the observed in vivo findings. Expression level of Gli1 was too low to be reliably detected in culture, likely due to absence of HH signaling which in vivo is provided exclusively by the lung epithelium.

The signaling of FGF10 and the expression of *Wnt5a* and *Foxd1* were blocked in culture with the FGFR inhibitor (Infigratinib at 1uM as in Manchado et al., 2016; Nakamura et al., 2015; Wong et al., 2018), *siWnt5a* (at 20nM as in Nemoto et al., 2012; Sakisaka et al., 2015; Zhao et al., 2017) and *siFoxd1*(at 20nM as in Li et al., 2021; Nakayama et al., 2015; Wu et al., 2018) respectively, and their effects on other SCMF genes were quantified by RT-PCR (Fig.4C). Identified genes with significant change are designated as targets of the gene/pathway perturbed, and their connections are plotted on the network (Fig.4D).

The network constructed on the new perturbation data has built connections between *Wnt5a* and certain fibroblast effector genes (i.e. *Acta2, Eln*) confirming its regulatory control on fibroblast growth and development. This finding in regard to this aspect of Wnt5a’s function resonates well with the clinical data from the study of some IPF patients (Martin-Medina et al., 2018). The architecture of the new construct reveals the identity of 3 transcription factors FOXD1, TBX2, and SOX8 forming a presumed connection hub. They are targeted by all 3 signaling pathways within the network and with each other as demonstrated by FOXD1’s regulation of *Tbx2* and *Sox8*, likely representing a core network subcircuit (Peter and Davidson, 2009) tasked to lockdown a regulatory state so that the signaling effect from the upstream can be stabilized.

### Alveolar epithelial genes are affected through WNT5a and FGF10 signaling

As mentioned above, both AT1 and AT2 cells were reduced in *Gcre*;*Igf1r-f/f* mutant lungs (Fig.2C). To identify any genetic alterations within these cells, we analyzed our FACS-sorted Tomato+ cells (Fig.3A), in which the lung epithelial cells reside, for the expression of a broad list of known AT1/AT2 signature genes, including their canonical markers and the ones identified by scRNAseq (i.e. Treutlein et al., 2014; Nabhan et al., 2018; LungMAP). The comparison between the control and the mutant data identified the genes as significantly altered (Fig.5A).

**Figure 5:**
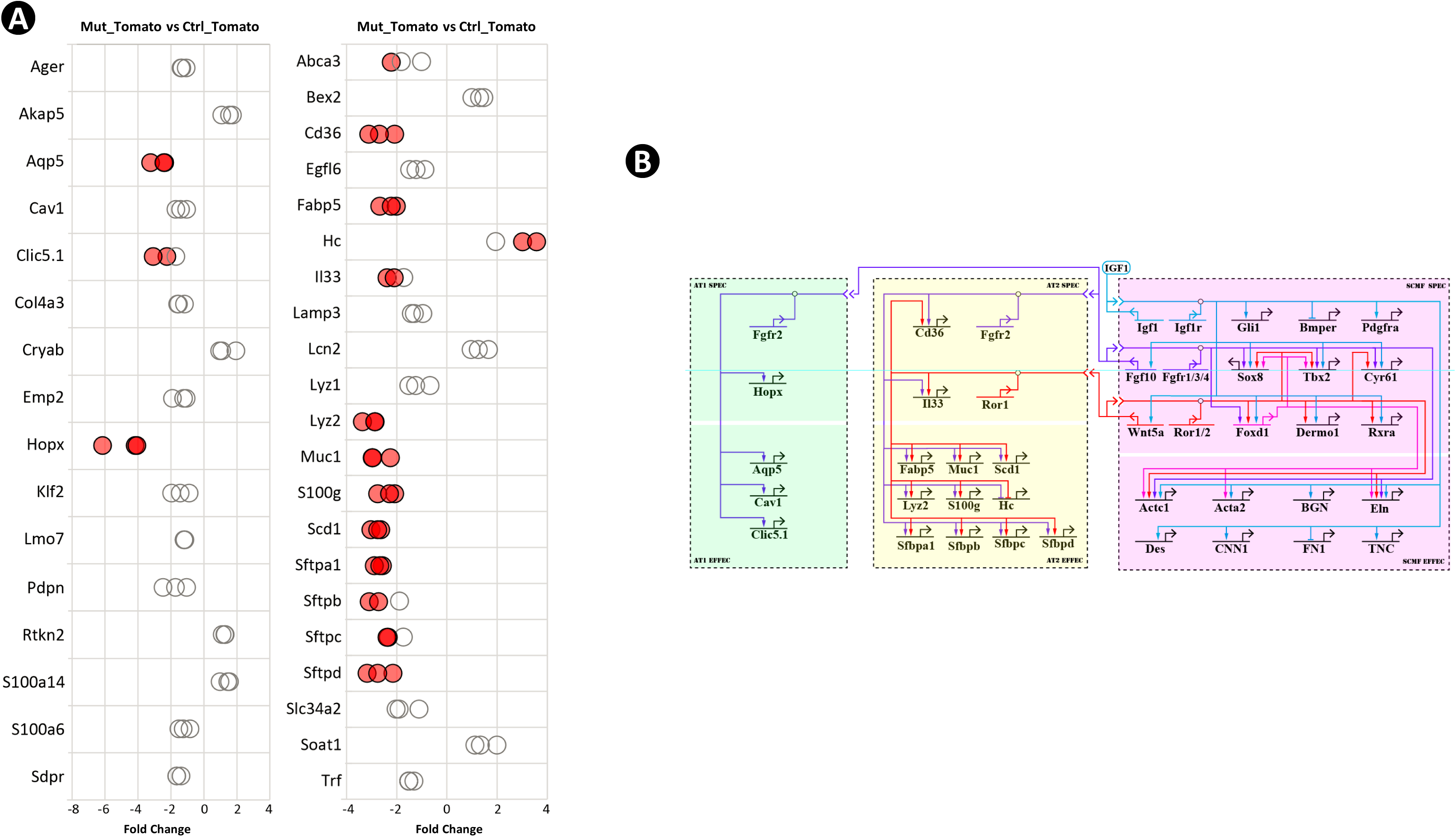
Epithelial genes affected in *Gli1-cre^ERT2^;Igf1r-f/f* mutant lung and their connections to IGF1 signaling from SCMF. A: RT-PCR data from selected AT1 and AT2 genes revealing their alteration in the mutant lung. B: The IGF1 signaling GRN during alveologenesis consisting of genes downstream of IGF1 signaling from three cell types (SCMF, AT2 and AT1) and the intracellular and intercellular regulatory connections among them.

Since our *Gcre;Igf1r-f/f* mutant was a SCMF targeted deletion, these changes must be a secondary consequence of intercellular cross communication. Within the altered SCMF gene list there are a total of 3 ligands: *Wnt5a*, *Fgf10,* and *Cyr61* (Fig.4D). CYR61 is more commonly recognized as a matricellular protein (i.e. Lau, 2011) rather than a classical growth factor such as FGF10 and WNT5A. Secretome-Receptome analysis indicates that FGF10 and WNT5a from SCMF communicate with alveolar epithelial cells through their cognate receptors, FGFR2 and ROR1 (Fig.4B, Figure 4-Figure Supplement 1).

The intercellular connections discovered above, plus the affected epithelial genes, were also added to the network as shown in Fig.5B. Multiple evidence has shown that IGF1 signaling can promote lung epithelial growth and development (Ghosh et al., 2013; Narasaraju et al., 2006; Wang et al., 2018b). Our work here reveals a nearly comprehensive look at the genetic pathway behind that.

### WNT5a is required for alveolar formation as inferred by the IGF1 signaling GRN

Our effort so far has led to construction of the IGF1 signaling GRN during alveologenesis. The network offers firstly a bird’s-eye view of how the genes involved in IGF1 signaling are connected in the form of a molecular circuitry for alveologenesis, and then a mechanistic perception of how this circuitry is initially turned on in SCMF for the function of these cells and then advances to the neighboring alveolar epithelial cells to influence their cellular activities.

Under this mechanistic view, the position where a gene is located on the architecture of the network and the connections it has with other genes manifests its role and impact on the network’s operation and outcome, hence its specific biological function in the process of alveolar formation. The network makes it possible to assess and predict a gene’s function before it is tested in vivo.

Mainly derived from our in vitro data, it was discovered WNT5a signaling on the current network is located immediately downstream of IGF1 and the signaling has connections with many IGF1 downstream genes (Fig.5B). This indicates WNT5A’s feed-forward role on the transduction of IGF1 signaling and its biological effects. Indeed, when *Wnt5a* was inactivated postnatally in mice, the mutant lungs exhibited a BPD-like phenotype with arrested alveologenesis, similar to what was seen in the *Igf1r* mutant (Figure 5-Figure Supplement 1; Li *et al.,* 2020). Inversely, the in vivo perturbation data collected from this *Wnt5a* mutant mouse model can be used to further define the regulatory connections where WNT5A is involved on the current IGF1 GRN.

### A GRN of similar components and wiring underlies human BPD

The expression of the mouse IGF1 signaling GRN regulatory genes in SCMF was also examined in lung samples from postmortem human BPD samples (Figure 6-Source Data 1). The regulatory genes, on the upper hierarchy of the network, determine the network’s outcome. In comparison to non-BPD lungs, 9 of the 12 genes examined were altered and 8 were altered in the same direction as they were defined in the mouse GRN (Fig.6). These findings indicate a genetic program of similar components and wiring underlying in human BPD.

**Figure 6:**
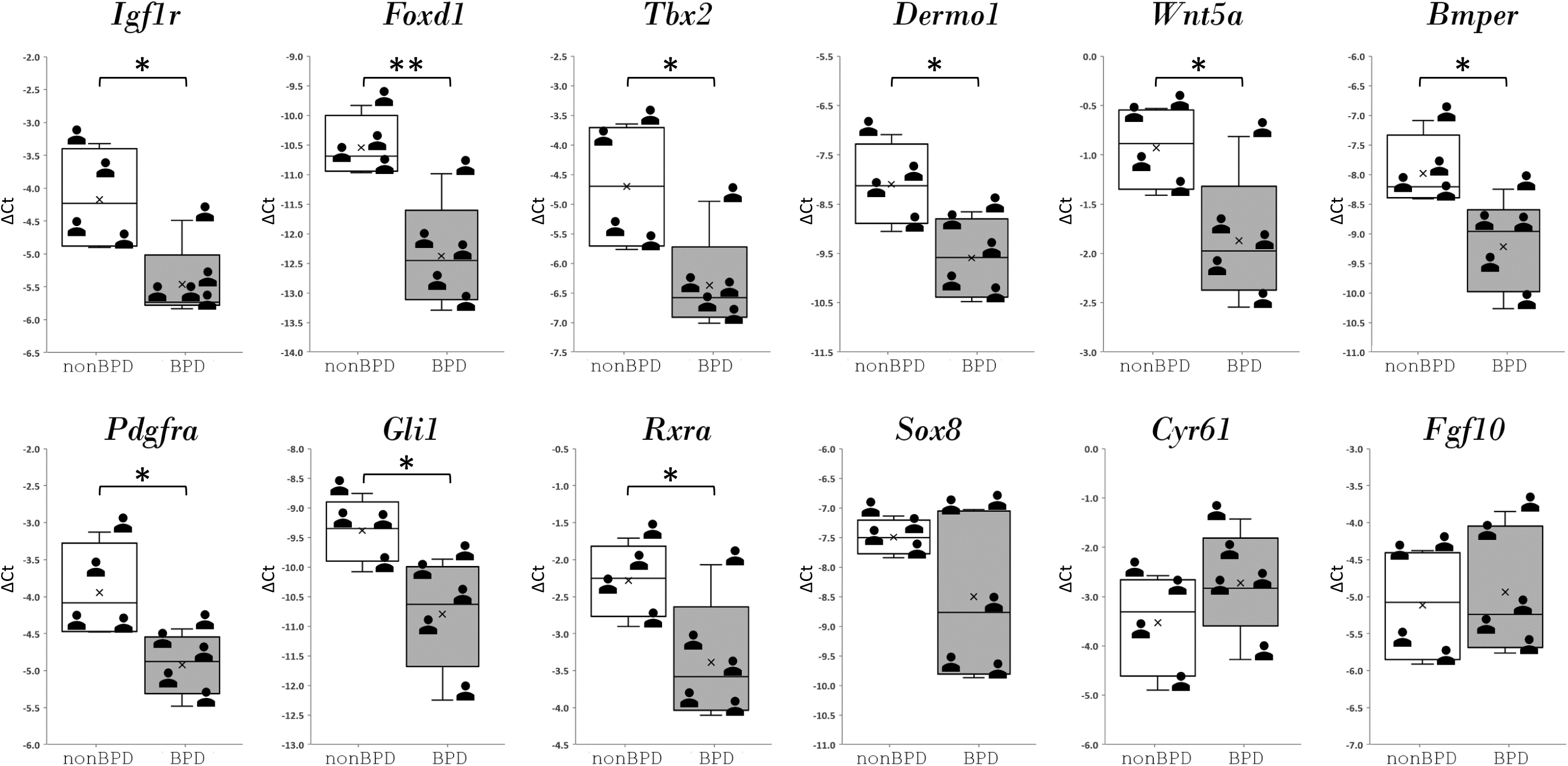
Regulatory genes from IGF1 signaling GRN and their expression in human bronchopulmonary dysplasia (BPD) lungs.

## Discussion

With massive gene expression data available (e.g., the LungMAP database) in this postgenomic era, one pressing task is to identify the function of genes, in particular, how they interact with one another (Przybyla and Gilbert, 2021). With gene regulatory network analysis as our targeted approach, we have constructed the IGF1 signaling GRN underlying alveologenesis using a mouse model of BPD.

The application of the GRN approach in the lung field is novel as it hasn’t been previously reported. Although not all the genes on the network have been examined by perturbation and the links haven’t been determined as direct or indirect, the regulatory connections running along the constructed GRN are already able to provide potential mechanistic explanation as to how the effect of IGF1 signaling is transduced from one gene to a constellation of its downstream genes, and from one cell type to another. Indeed, blocking WNT5A signaling is confirmed to produce a mouse BPD-like phenotype that mimics Igf1r-/- lungs as inferred by the network (Figure 5-Figure Supplement 1; Li *et al.,* 2020). Signaling by IGF1, FGF10 and WNT5A have long been recognized in having roles in alveologenesis (i.e. Chao et al., 2016; He et al., 2021; Li et al., 2020; Zhang et al., 2020). The present GRN reveals a full genetic program and the cross-talk among them. The fact that all these signaling pathways have connections on the specification and effector genes of both alveolar mesenchymal and epithelial cells provides a direct causal link between the signaling and alveolar development.

Behind the overall process of alveologenesis is a much larger signaling gene regulatory network where several different cell types are involved. Within the mesenchymal cell type only, it is known that blocking IGF1, WNT5a, PDGFa, and TGFb signaling leads to impairment in development of alveoli, though with varying severities (Gao et al., 2022; He et al., 2021; Li et al., 2020; Li et al., 2019; Zhang et al., 2020). The sum of these leads to the construction of the hierarchical regulatory connections of these pathways within SCMF (Fig.7). Once signaling pathways from alveolar epithelium, endothelium, and immune cells are included, a much larger and more comprehensive signaling GRN behind alveologenesis is expected to emerge.

**Figure 7:**
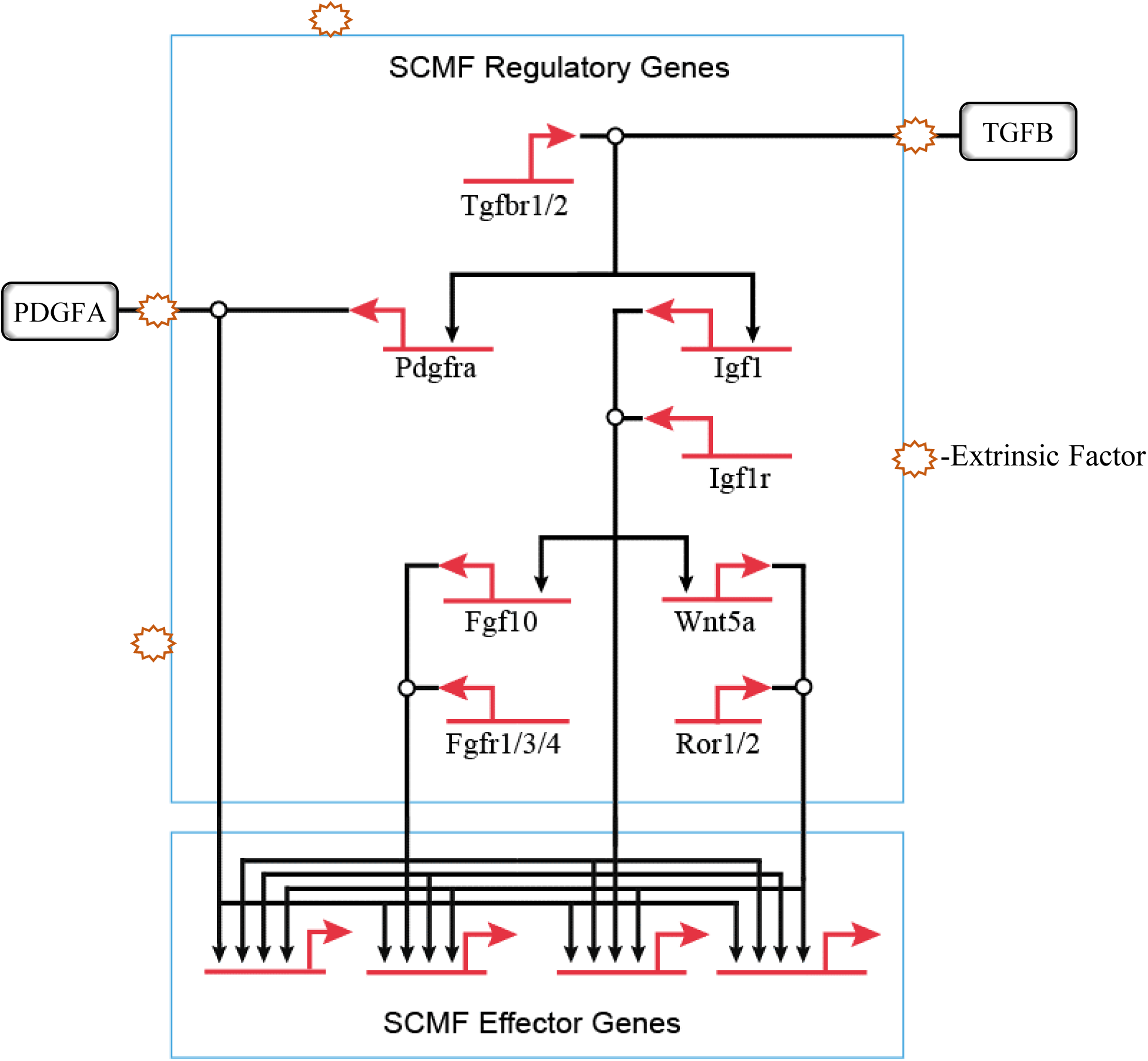
The hierarchical connections among the signaling pathways within SCMF during alveologenesis.

Clinically, BPD is caused predominantly by extrinsic factors which in the first place would interfere with normal cell’s extracellular activities including cell signaling and communication. From the GRN perspective, BPD is a developmental disease when the signaling GRN for alveologenesis is derailed and disrupted by these extrinsic factors (Fig.7).

The type and effect of these extrinsic factors must be taken into consideration when studying this disease. There was a reported increase in Wnt5A expression by mesenchymal cells from hyperoxia BPD models (Sucre et al., 2020)—a finding seemingly contradictory to our data on Wnt5a. The etiology of BPD is multifactorial in that it involves a plethora of factors such as lung immaturity, injury, inflammation, and genetic defects. In the injury model, the two opposite conditions, hyperoxia and hypoxia, can both cause lung injury and lead to BPD-like phenotypes. In our model, the genetic defect from the targeted genetic manipulation is the culprit behind the lungs of the BPD phenotype. It is expected that genes on the network may respond differently to these different conditions and challenges. The derailed gene expression on the network, either increased or decreased, can both disrupt the signaling GRN for alveologenesis and cause BPD.

The aforementioned network disruption may occur within different cell types, onto different signaling pathways, and upon different hierarchy of the network (Fig.7). From the network’s structural view, the disruption at the higher hierarchy has greater manifestation, thus leading to a higher severity of disease. TGFB signaling, atop the whole hierarchy of the signaling GRN on Fig.7, exhibits in its mutant mouse model the most severe BPD-like phenotype when compared to mutants from the other signaling pathways under it (Chao et al., 2017; Gao et al., 2022; He et al., 2021; Li et al., 2020; Li et al., 2019). Also, different signaling sitting at various hierarchical levels has different coverage (sum of genes under the signaling’s control) on the network. If disruption occurred beyond a signaling’s coverage, the treatment by targeting this signaling would be off target. IGF1 signaling is found to be in the middle hierarchical level of the constructed network (Fig.7). By targeting simply one such signaling and hoping to treat BPD altogether will surely not always work.

Our GRN view on alveologenesis presents a transformative viewing of BPD and could potentially help in designing novel strategies to prevent and treat it. As for prevention, the GRN suggests a study centered on the network’s periphery with its focus on cutting off the extrinsic factors and blocking their connections to the intrinsic alveologenesis GRN. For treatment, it is a study on the network itself focusing on recovering and rescuing the signaling pathways that have been disrupted. However, all this relies on the foundational construction of the whole alveologenesis GRN—which includes all major signaling pathways involved within and between each cell type and the network assembly of the hierarchical regulatory connections among them. Though the network construction is currently not experimentally accessible in human models, our work shows the alveologenesis GRN is well conserved between mouse and human lungs. With the mouse model, the whole signaling GRN behind alveologenesis can be decoded as we did in this paper. The impact of extrinsic factors/signaling pathways and their manipulation on the network can be modeled and tested as well. The insights collected can then be used to guide health delivery for BPD clinically. The general process would then be as follows: clinical patient exams and tests are used to determine the extrinsic factors; cell type gene expression data from bronchoalveolar lavage and/or biopsy are used to reveal genes altered within the alveolar compartments in patient lungs; implied extrinsic factors and altered genes are mapped onto the alveologenesis GRN where points of network disruptions can thus be defined; therapies proven successful from mouse modeling data can then be clinically pursued to recover/rescue the disruption points in the patient.

The GRN is traditionally a network built solely on chemical interactions (RNA, DNA, protein and other chemicals). Having said that however, it has been recognized that these interactions can also happen at mechanical and electrical level (i.e. Azeloglu and Iyengar, 2015). Mechanosignaling is of particular interest in the lung as the organ itself is born to function through its non-stopping mechanical movement. New technologies, including whole genome CRISPR perturbation and screening, epigenomic profiling, high resolution ChIP-seq, spatial genomics and cis-regulatory analysis, are sprouting with their potential use for a high throughput GRN construction (i.e. Cusanovich et al., 2018; Eng et al., 2019; Sanson et al., 2018; Skene and Henikoff, 2015). Our journey to decode the alveologenesis GRN is also an adventure to construct the new generation GRNs with rising technologies.

A web resource of the network and the data presented in this paper is made available to the public: https://sites.google.com/view/the-alveologenesis-grn/home, which we hope can be used as an online repository to promote any further studies and collaborations in this direction.

## MATERIALS AND METHODS

### KEY RESOURCES TABLE

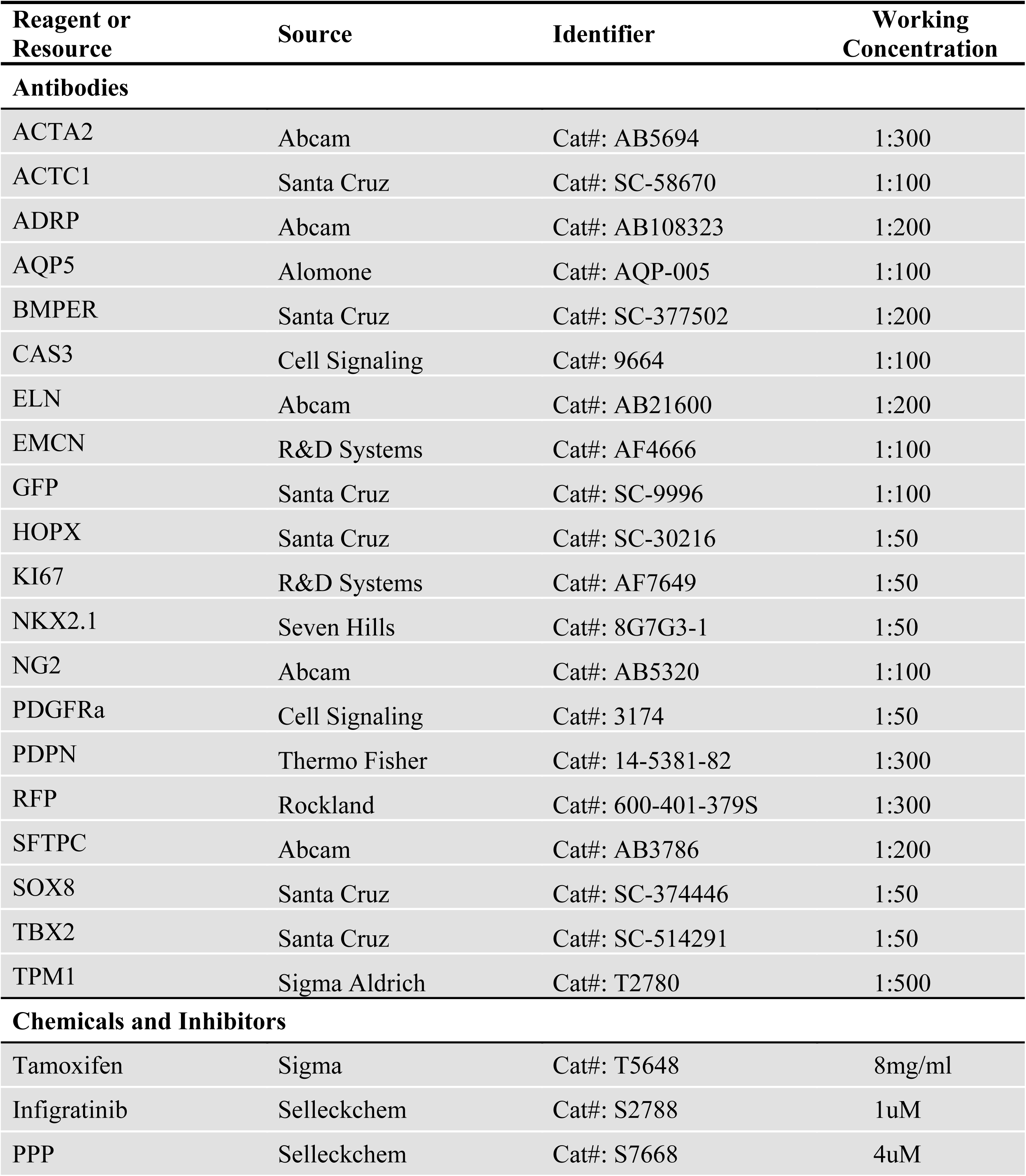
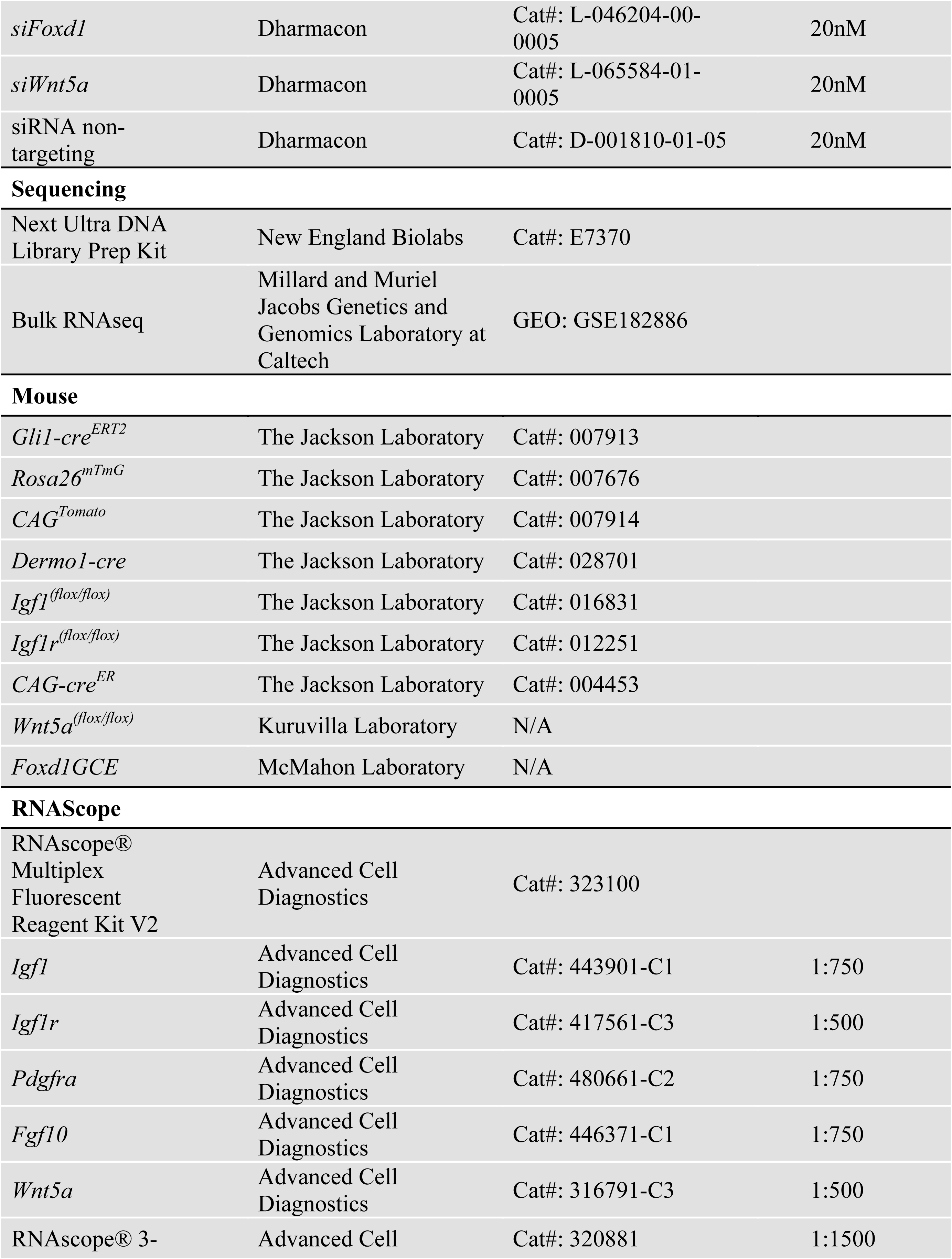
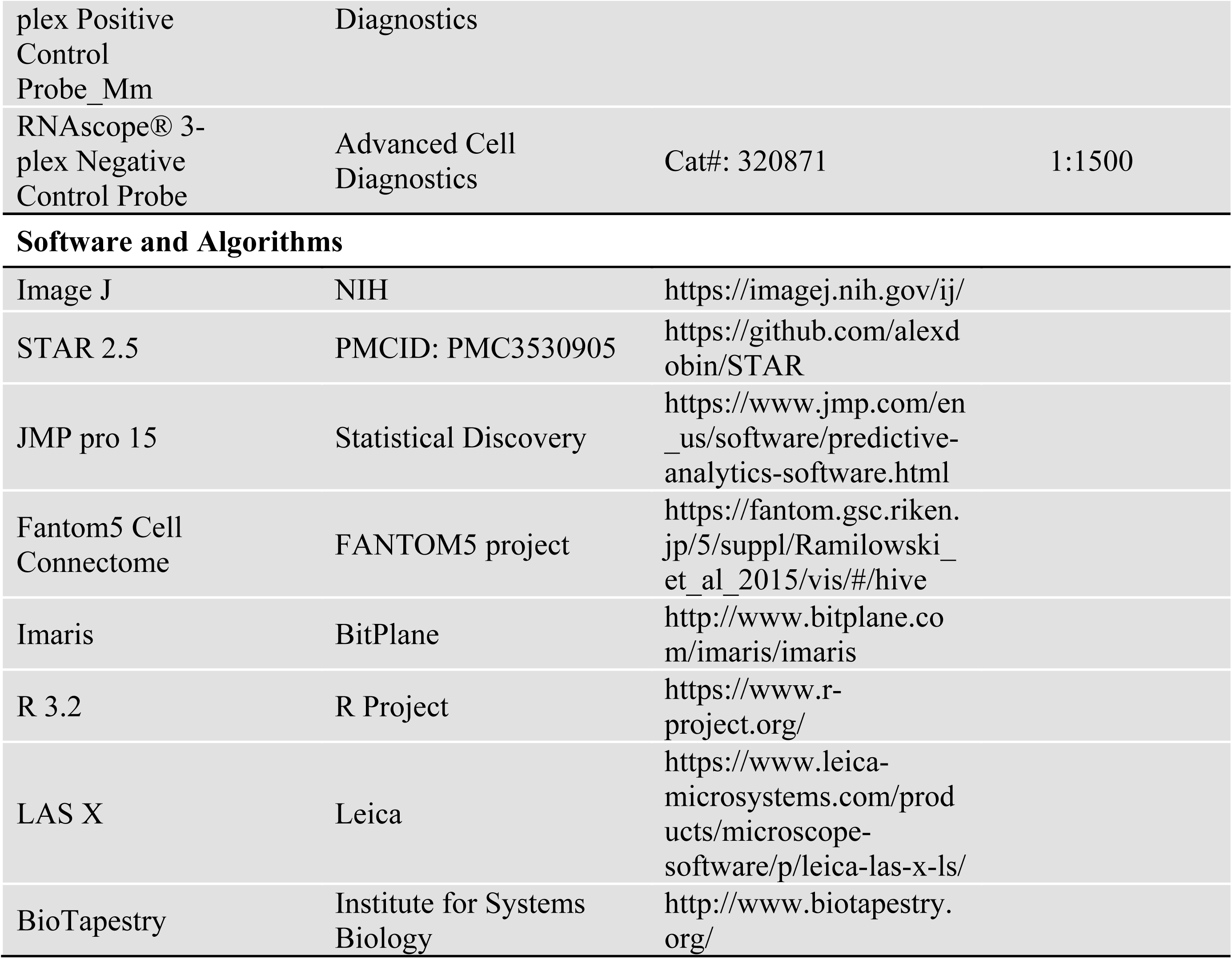

#### 1. Mouse Breeding and Genotyping

All animal studies were conducted strictly according to protocols approved by the USC Institutional Animal Care and Use Committee (IACUC) (Los Angeles, CA, USA). The mice were housed and maintained in pathogen-free conditions at constant room temperature (20–22 °C), with a 12 h light/dark cycle, and free access to water and food. *Dermo1-cre*, *Gli1-cre^ERT2^, Rosa26^mTmG^, CAG^tdTomato^, Igf1^(flox/flox)^, Igf1r^(flox/flox)^*, and *CAG-cre^ER^* mice were purchased from the Jackson Laboratory. *Foxd1GCE* mice were generated by McMahon lab at USC. *SPC^GFP^* mice were generated by Wright lab at Duke University. Wnt5a*^(flox/flox)^* mice were provided by Kuruvilla lab at Johns Hopkins University.

*Dermo1-cre;Igf1^(flox/flox)^ (Dcre;Igf1-f/f)* and *Dermo1-cre;Igf1r^(flox/flox)^ (Dcre;Igf1r-f/f)* mice were generated by breeding *Dermo1-cre with Igf1^(flox/flox)^* and *Igf1r^(flox/flox)^* respectively.

*Gli1-cre^ERT2^;Rosa26^mTmG^ mice* were generated by breeding *Gli1-cre^ERT2^* and *Rosa26^mTmG^* mice.

*Gli1-cre^ERT2^;Igf1r^(fox/flox)^* (*Gcre*;*Igf1r-f/f*) mice were generated by breeding *Gli1-cre^ERT2^* mice with the *Igf1r^(fox/flox)^* mice.

*Rosa26^mTmG^;Igf1r^(fox/flox)^ (mTmG;Igf1r-f/f)* mice were generated by breeding *Rosa26^mTmG^* mice with the *Igf1r^(fox/flox)^* mice.

*Gli1-cre^ERT2^;Rosa26^mTmG^;Igf1r^(fox/flox)^ (Gcre;mTmG;Igf1r-f/f)* mice were generated by breeding *Gli1-cre^ERT2^;Igf1r^(fox/flox)^* mice with the *Rosa26^mTmG^;Igf1r^(fox/flox)^* mice.

*Foxd1GCE;CAG^tdTomato^(Fcre;Tomato)* mice were generated by breeding *Foxd1GCE* mice with the *CAG^tdTomato^* mice.

*CAG-cre^ER^;Wnt5a^(fox/flox)^ (Ccre;Wnt5a-f/f)* mice were generated by breeding *CAG-cre^ER^* mice with the *Wnt5a^(fox/flox)^* mice.

Genotyping of the transgenic mice was performed by PCR with genomic DNA isolated from mouse tails. The forward (F) and reverse primers (R) for transgenic mouse genotyping are listed in Figure 3-Source Data 2.

#### 2. Tamoxifen Administration

A single dose of Tamoxifen (8mg/mL in peanut oil) was administered by oral gavage to neonates at postnatal day 2 (PN2, 100μg per pup) with a plastic feeding needle (Instech Laboratories, PA). Neonatal lungs were collected between PN7 and PN30 for morphological, immunohistochemical, cellular, and molecular biological analyses.

#### 3. Mouse lung tissue

Mice were euthanized by CO2 inhalation at the time of tissue harvest. Chest cavity was exposed and lungs cleared of blood by perfusion with cold PBS via the right ventricle. Lungs were inflated with 4% formaldehyde under constant 30cm H2O pressure and allowed to fix overnight at 4°C. Tissue was dehydrated through a series of ethanol washes after which they were embedded in paraffin and sectioned.

#### 4. Immunohistochemistry

H&E staining was performed as usual, and morphometric measurements were made using ImageJ. Immunofluorescent staining was performed as previously described using paraffin-embedded lung sections (Li et al., 2019). In brief, five micrometer (*μm*) tissue sections were deparaffinized, rehydrated, and subjected to antigen retrieval. After blocking with normal serum, the sections were probed with primary antibodies at 4° C overnight. Combinations of Alexa Fluor Plus secondary antibodies (Thermo Fisher Scientific) were applied for fluorescent detection of above specific primary antibodies. Nuclei were counterstained with 4′,6-diamidino-2-phenylindole (DAPI). Primary antibodies used and their sources are listed in the Key Resources Table below. Images were made with Leica DMi8 fluorescence microscope and processed with Leica LAS X and ImageJ.

#### 5. RNAScope

Samples were fixed in 10% neutral buffered formalin, dehydrated with ethanol, and embedded in paraffin wax. 5*μm* Sections from paraffin blocks were processed using standard pretreatment conditions per the RNAscope multiplex fluorescent reagent kit version 2 (Advanced Cell Diagnostics) assay protocol. TSA-plus fluorescein, Cy3 and Cy5 fluorophores were used at different dilutions optimized for each probe. RNAScope probes used are listed in the Key Resources Table below. Images were made with Leica DMi8 fluorescence microscope and processed with Leica LAS X and ImageJ.

#### 6. Mouse lung single-cell dissociation

Single-cell suspension was prepared as described in Adam et al. (2017) with all the procedures performed on ice or in cold room. Mice were euthanized and the lungs were perfused with PBS as described above. The lungs were inflated with cold active protease solution (5mM CaCl2, 10 mg/ml Bacillus Licheniformis protease), dissected, and transferred to a petri dish where the heart, thymus, and trachea were removed. The lobes were minced using a razor blade. The minced tissue was then immersed in extra cold active protease solution for 10 min and triturated using a 1ml pipette. This Homogenate was transferred to a Miltenyi C-tube with 5ml HBSS/DNase (Hank’s Balanced Salt Buffer) and the Miltenyi gentleMACS lung program was run twice on GentleMACs Dissociator. Subsequently, this suspension was passed through a 100um strainer, pelleted at 300g for 6 minutes, suspended in 2ml RBC (Red Blood Cell) Lysis Buffer (BioLegend), and incubated for 2 min. At this point, 8ml HBSS was added and centrifuged again. The pellet was suspended in HBSS, then filtered through a 30um strainer. The suspension was pelleted again and finally suspended in MACS separation buffer (Miltenyi Biotec) with 2% FBS for Fluorescence-activated cell sorting (FACS). Cell separation and viability were examined under the microscope and through Vi-CELL Cell Counter after staining with Trypan blue.

#### 7. Flow Cytometry and Cell Sorting

FACS was performed on a BD FACS Aria II at stem cell Flow Cytometry Core at the Keck School of Medicine of USC. The sorting was gated on viability, singlets, GFP and/or Tomato. GFP+ and/or Tomato+ cells were collected as needed. For cell culture, cells were sorted in DMEM containing 10% FBS. For RNA, cells of interest were collected in Trizol-LS reagent (ThermoFisher).

#### 8. Bulk RNA-seq and scRNAseq data analyses

Lungs dissected from Sftpc^GFP^ mouse at PN14 were dissociated into single cell suspension and GFP+ cells were sorted and collected as described above. RNA was extracted using Qiagen RNeasy Micro kit and then submitted to the Millard and Muriel Jacobs Genetics and Genomics Laboratory at Caltech for sequencing, which was run at 50bp, single end, and 30 million reading depth. The unaligned raw reads from aforementioned sequencing were processed on the PartekFlow platform. In brief, read alignment, and gene annotation and quantification, were based on mouse genome (mm10) and transcriptome (GENECODE genes-release 7). Tophat2 and Upper Quartile algorithms were used for mapping and normalization. The RNA-seq data have been deposited with GEO under the accession number GSE182886.

The raw scRNAseq datasets (GSE160876 and GSE165063 from Negretti et al., 2021, GSE149563 from Zepp et al., 2021) were downloaded from the SRA server. The 10x Genomics’ Cell Ranger pipeline was used to demultiplex raw base call (BCL) files into FASTQ files, perform alignment, filtering, barcode counting, and UMI counting, combine and normalize counts from multiple samples, generate feature-barcode matrices, run the dimensionality reduction, clustering, and gene expression analysis using parameters once they were provided in the original papers. Quality control was done additionally in Partek Flow to filter out cells with excess mitochondrial reads and possible doublets and remove batch effects. Loupe Browser was used for data visualization and analysis including cell clustering, cell counting, cell type classification, gene expression and comparative analysis.

#### 9. Neonatal lung myofibroblast culture and treatment

FACS sorted GFP+ cells from PN5 neonatal lungs of *Gli1-cre^ERT2^;Rosa26^mTmG^* mice were suspended in DMEM containing 10% FBS, plated in 24-well culture plates at 50,000 cells/well, and incubated at 37°C with 5% CO2 on day 1 after sorting. On day 2 after sorting, the attached myofibroblasts were washed with PBS and cultured in fresh medium with inhibitors or siRNAs as indicated in each experiment. The ON-TARGETplus SMARTpool siRNAs from Dharmacon were used, and the transfection was done using DharmaFECT™ Transfection Reagents. On day 4, the cells were collected for RNA analyses. The cells were authenticated for absence of contaminations.

#### 10. Real-time Quantitative Polymerase Chain Reaction (real-time RT-PCR)

Neonatal mouse lung cells were collected from FACS or cell culture as described above. The RNA was isolated with Direct-zol™ RNA MiniPrep kit according to the manufacturer’s protocol (ZYMO Research). Following RNA purification, cDNA was generated using the SuperScript IV First-Strand Synthesis System (ThermoFisher). Expression of selected genes was quantified by Quantitative Real Time RT-PCR performed on a Light Cycler (Roche) or 7900HT fast real-time PCR system (Applied Biosystems) using SYBR green reagents (ThermoFisher). The deltaCT method was used to calculate relative ratios of a target gene mRNA in mutant lungs compared to littermate control lungs. *Gapdh* was used as the reference gene. Primers for each gene were designed on IDT website and the specificity of their amplification was verified by their melting curve. Sequences of the primers are listed in Figure 3-Source Data 2.

#### 11. Human neonatal lung samples

BPD and non-BPD postnatal human lung tissues were provided by the International Institute for the Advancement of Medicine and the National Disease Research Interchange, and were classified exempt from human subject regulations per the University of Rochester Research Subjects Review Board protocol (RSRB00056775).

#### 12. Secretome-Receptome analyses

Cell to cell communications were predicted using a published Fantom5 Cell Connectome dataset linking ligands to their receptors (STAR Methods) (Ramilowski et al., 2015). The ligands and receptors were identified from the following bulk RNAseq datasets: GSE126457 for SCMF (Li et al., 2019), GSE182886 for AT2 (submitted with this paper), GSE106960 for AT1 (Wang et al., 2018a), and the following scRNAseq datasets: GSE160876 (Schuler et al., 2021), GSE165063 (Negretti et al., 2021), and GSE149563 (Zepp et al., 2021). The IGF1 signaling GRN was drawn in BioTapestry software developed by Longabaugh et al., 2012.

#### 13. Quantification and Statistical Analysis

In gene expression quantification using RT-PCR, at least three biological replicates (in different cases including lungs/FACS sorted cells/cultured cells) for each experimental group (Ctrl vs Mut, FACS sorted cell lineage #1vs #2, treated vs untreated, BPD vs nonBPD) were used. Measurement for each biological replicate was repeated three times. The Ct (cycle threshold) was normalized to *Gapdh*, and the final result was presented as deltaCt or fold change. In morphometric quantification and cell counting, four lungs for each experimental group (Ctrl vs Mut) were used. Left lobe and right inferior lobe from each lung were targeted. Five images from each lobe after staining were analyzed for morphometric quantification (at 10x magnification) and cell counting (at 40x magnification). A two-tailed Student’s t-test was used for the comparison between two experimental groups and a one-way ANOVA was used for multiple comparisons. Quantitative data are presented as mean values +/-SD. Data were considered significant if p < 0.05.

## Acknowledgements

We thank Dr. Andrew P. McMahon (Keck School of Medicine of USC) for providing the *Foxd1GCE;CAG^tdTomato^* mice. We thank Dr. Gloria S. Pryhuber (University of Rochester Medical Center) for providing the human BPD samples. We thank Arnold Sipos (Keck School of Medicine of USC) for help with imaging and Sean Gao (Arcadia High School/Duke University) for data analysis, editing, and the construction of the Alveologenesis GRN website.

## Data Availability Statement

Additional data that support the findings of this study are available on request from the corresponding author.

---

**Figure 1-Figure Supplement 1:**
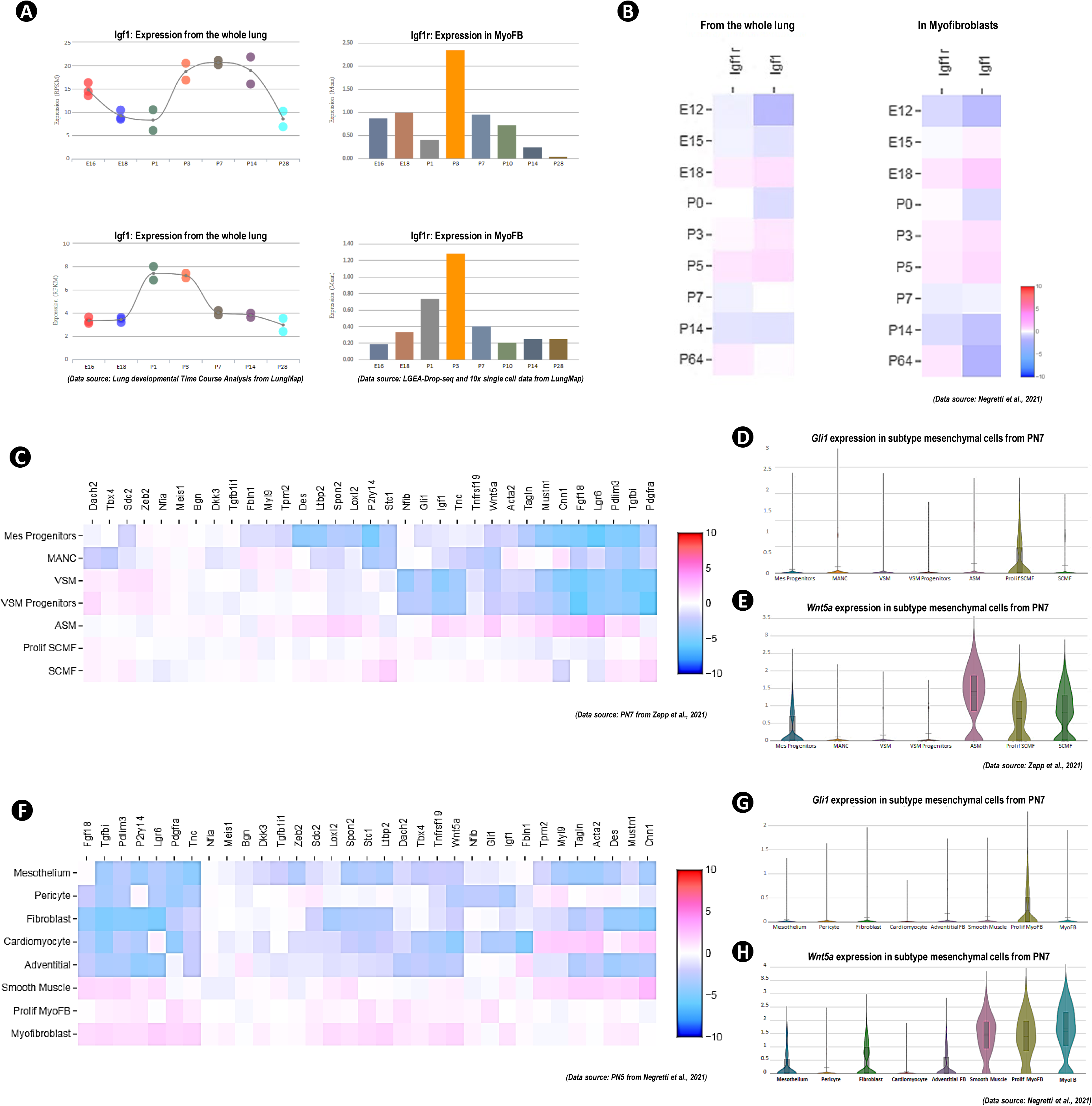
Igf1, Igf1r and SCMF/MyoFB marker gene’s expression in mouse lung from published public data sources. A, B: Expression profiling of Igf1 and Igf1r from whole lung and in SCMF/MyoFB cells during lung development as calculated on data from LungMap (A) and from Negretti et al., 2021 (B). C-H: Heatmap and violin plots of collected SCMF/MyoFB marker gene’s expression within mesenchymal sub cell clusters as calculated on data from Zepp et al., 2021 (C-E) and Negretti et al., 2021 (F-H).

**Figure 2-Figure Supplement 1:**
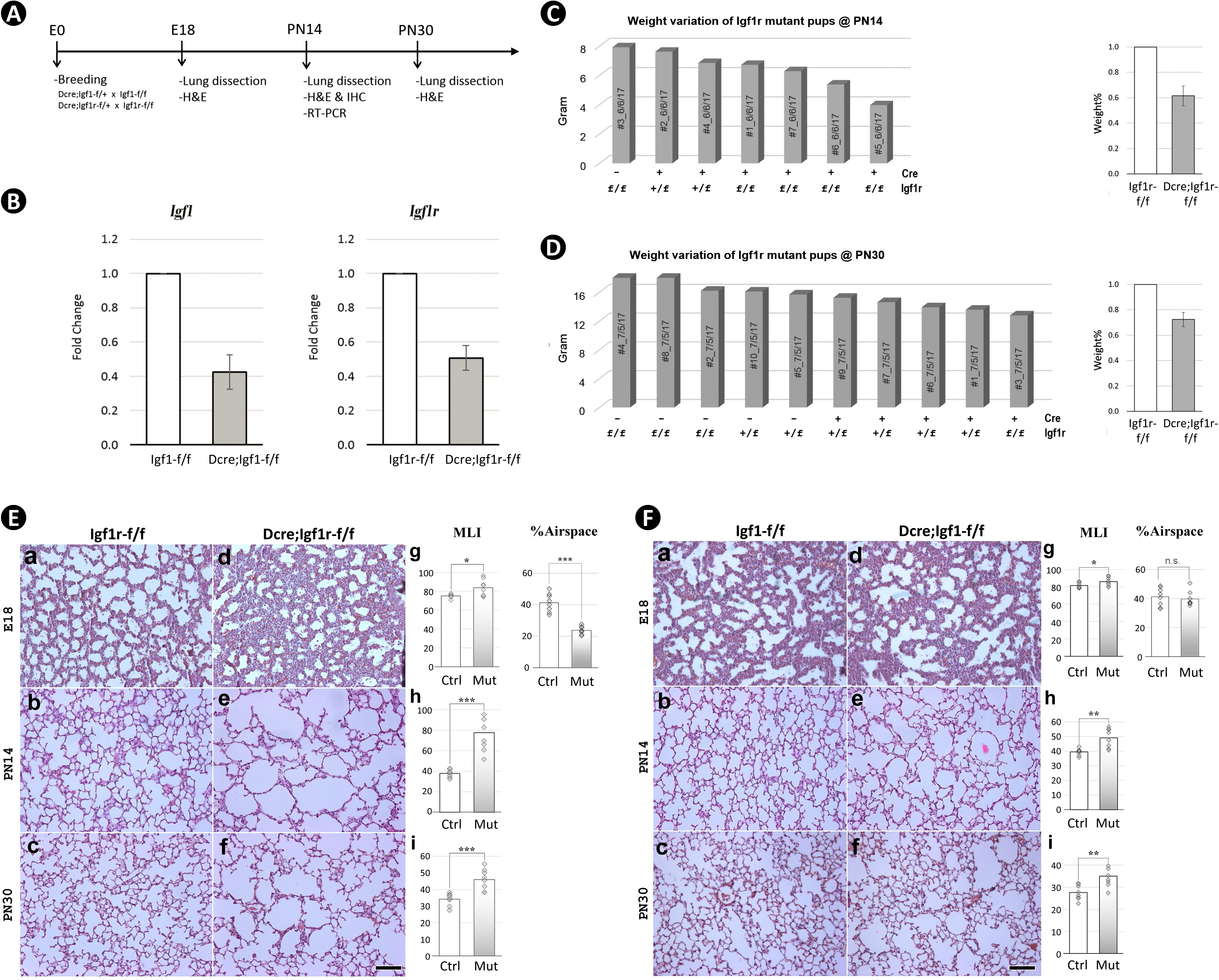
Mesoderm constitutive deletion of *Igf1* and *Igf1r* from mouse lung. A: Schematic of the experimental protocol. B: Impact of inactivation of *Igf1* (left) and *Igf1r* (right) by *Dermo1-cre* on their respective transcripts in PN14 lungs. Data is represented as mean +/- SD of three lungs from each experimental group. C: Body weight variation of pups at PN14 of one single litter from the breeding of *Dermo1-cre;Igf1r-f/-* and *Igf1r-f/f* (left graph). Body weight comparison of pups between the control and the homozygous mutant (right graph). Data is represented as mean +/- SD of 4 mice from each experimental group. D: Same experiment as done in C for pups at PN30. E: H&E staining of lung sections from *Igf1r* control (a-c) and *Dermo1-cre* mutant (d-f) mice and their morphometric measurements (MLI&%Airspace – see Figure 2-Figure Supplement 2E,F for more information) (g-i) at E18, PN14, and PN30. F: H&E staining of lung sections from *Igf1* control (a-c) and *Dermo1-cre* mutant (d-f) mice and their morphometric measurements (g-i) at E18, PN14, and PN30. Scale bar: 100um for all images. *p*-value: * stands for 0.05-0.001, ** for 0.01-0.001, *** for <0.001, n.s. for not significant. The same designation is used throughout the paper.

**Figure 2-Figure Supplement 2:**
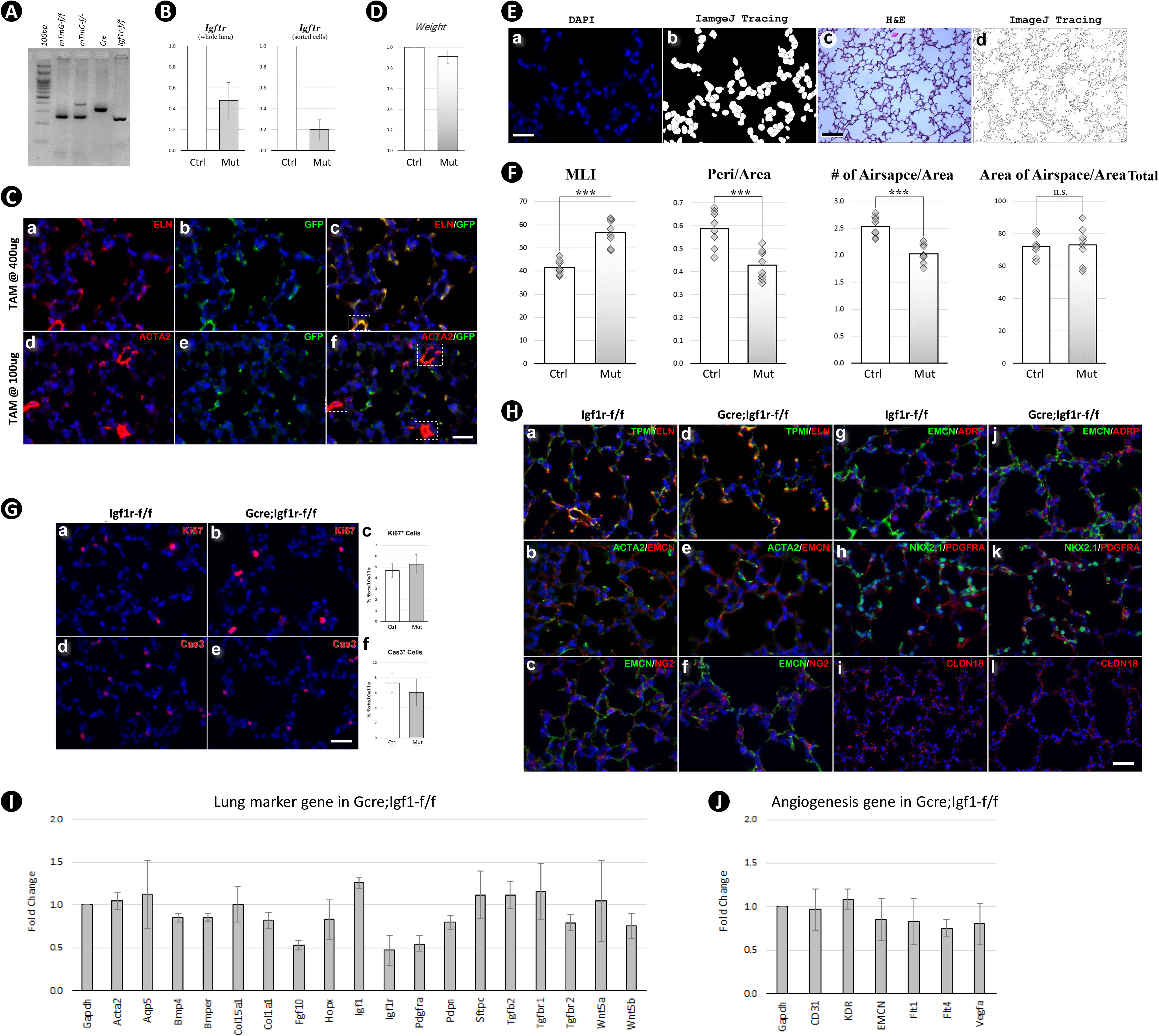
Postnatal inactivation of *Igf1r* in secondary crest myofibroblasts. A: genotyping of pups from the breeding between *Gli1-cre^ERT2^*;*Igf1r-f/f* and *Igf1r-f/f;mTmG* mice. B: *Igf1r* expression in control and mutant lungs at PN14 using whole lung (left) and FACS sorted GFP+ cells (right). Data is represented as mean +/- SD of three lungs from each experimental group. C: Immunostaining for Eln, Acta2, and GFP on PN14 lung sections from *Gli1-cre^ERT2^;mTmG* mice which were tamoxifen-treated at PN2 with 400ug (a-c) and 100ug (d-f) per pup. Smooth muscle fibers (outlined in c&f) were labeled by GFP at higher but not at lower TAM dosage. Scale bar: 25um for all images. D: Weight comparison between Ctrl and Mut at PN14. Data is represented as mean +/- SD of three mice from each experimental group. E: A sample of how to use ImageJ on cell counting (a,b) and morphometric measurements (c,d) where number of separated airspaces (*N*), perimeter (*Peri*) and area (*Area*) of each airspace can be calculated. Scale bar: 25um in a and 100um in c. F: Morphometric comparison between Ctrl and Mut at PN14 using different indexes. MLI is defined as in Crowley et al., 2019 and the others are quantified as below: 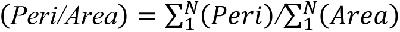, 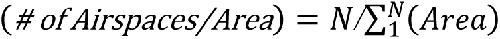, 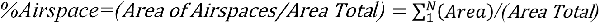. G: Immunostaining for Ki67 (a,b) and cleaved Caspase-3 (d,e) on control and mutant lungs at PN14. The percentage of cells identified from above staining was compared between the control and mutant (c,f). Data is represented as mean +/- SD of three lungs from each experimental group. Scale bar: 25um for all images. H: Immunostaining for mesenchymal marker ELN (a,d), TPM1 (a,d), ACTA2 (b,e), PDGFRA (h,k), epithelial marker NKX2.1 (h,k), CLDN18 (i,l), endothelial marker EMCN (b,c,e,f,g,j), lipofibroblast marker ADRP (g,j), and pericyte marker NG2 (c,f) using control and mutant lungs at PN14. Scale bar: 50um for i, l and 25um for all the other images. I: Comparative expression of selected lung marker genes between control and mutant lungs at PN14. Data is represented as mean +/- SD of three lungs from each experimental group. J: Comparative expression of angiogenesis related genes between control and mutant lungs at PN14. Data is represented as mean +/- SD of three lungs from each experimental group.

**Figure 3-Figure Supplement 1:**
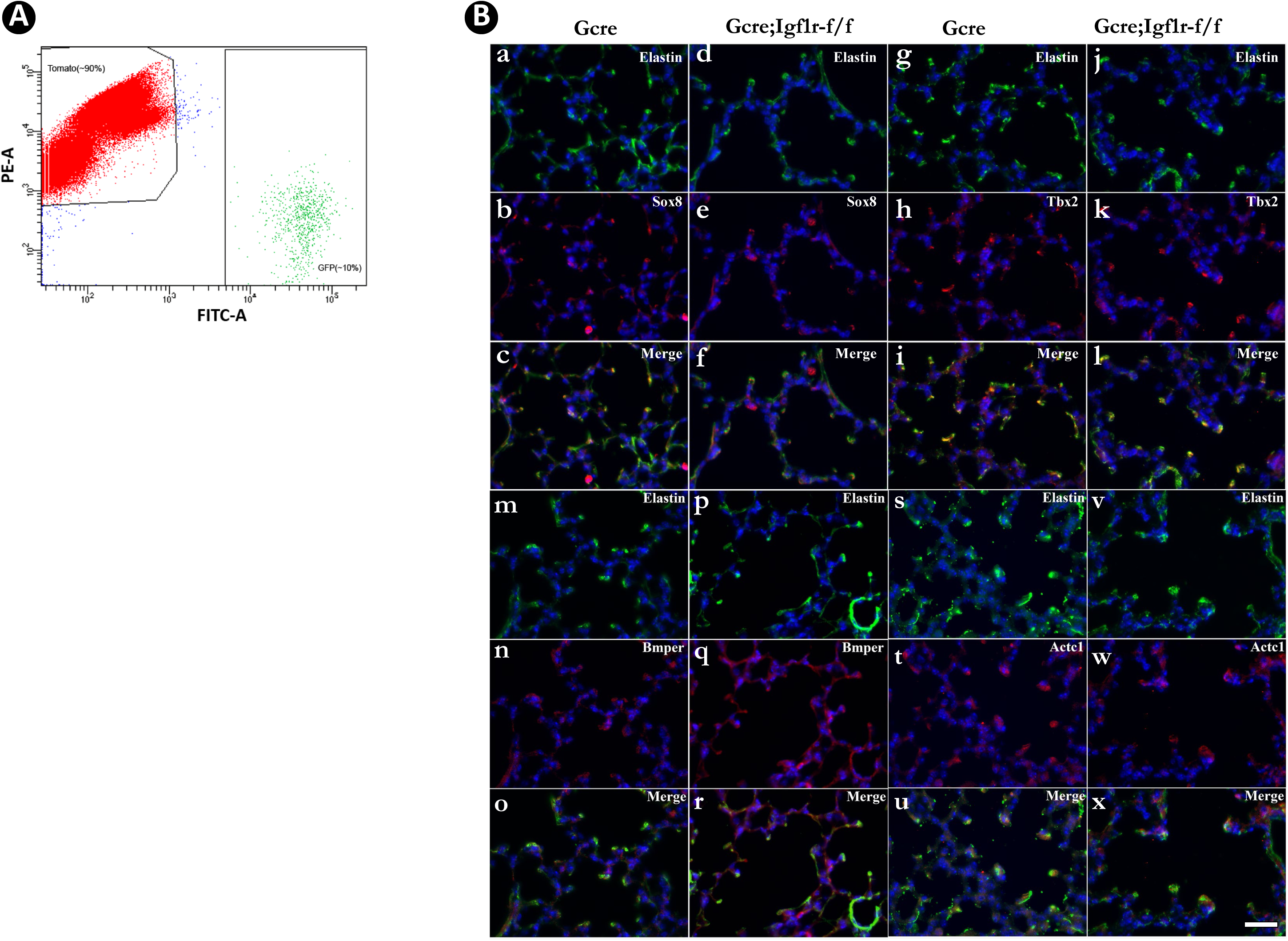
Identification of SCMF genes altered in *Gli1-cre^ERT2^;Igf1r-f/f* mutant lung. A: GFP+ and Tomato+ cells are separated by FACS. B: Spatial expression of SOX8/ELN (a-f), TBX2/ELN (g-l), BMPER/ELN (m-r), and ACTC1/ELN (s-x) in the alveolar compartment in control and mutant lungs at PN14 as detected by immunostaining. Scale bar: 25um for all images.

**Figure 3-Source Data 1: Table of genes with their RT-PCR data showing their level of expression in SCMF, enrichment in SCMF, and differential expression between control and mutant lungs, and annotation of their cellular expression in the lung based on LungMAP scRNAseq data.** Cells in red: data points which meet the cutoff criteria designated in the paper or the cell type where the gene is annotated as expressing.

**Figure 3-Source Data 2: List of the primers used in the paper.** Primers are for *Mus musculus* by default and for *Homo sapiens* when denoted with Hs.

**Figure 4-Figure Supplement 1:**
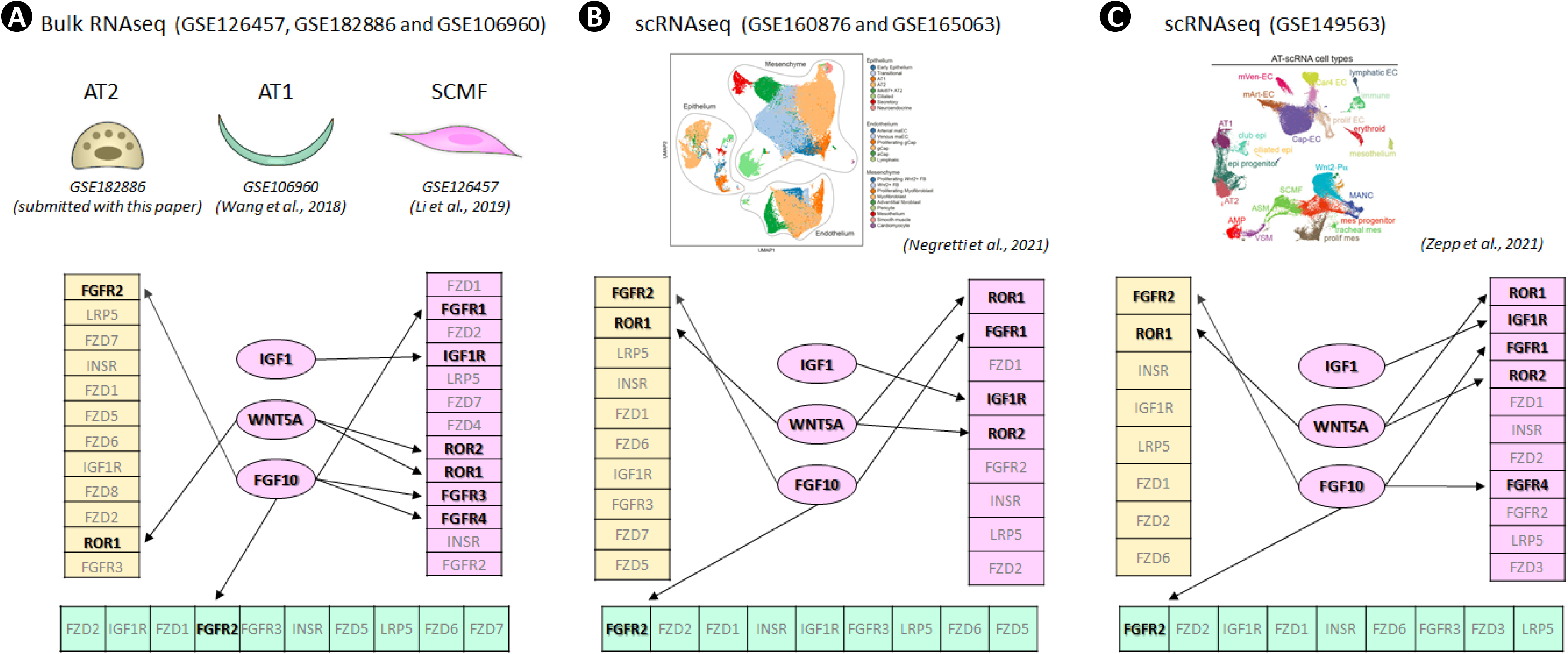
Secretome-receptome analysis of cellular communications between *IGF1*, *WNT5A* and *FGF10* ligands from SCMF and receptors from SCMF, AT2 and AT1 as calculated on different data sources: bulk RNAseq data (A), scRNAseq data from Negretti et al., 2021 (B) and scRNAseq data from Zepp et al., 2021 (C). Receptors are rendered as magenta for those from SCMF, yellow for AT2, and cyan for AT1; are arranged based on their abundance within each cell type with higher quantities on top/left and lower quantities on bottom/right. Ligands are connected to the receptors predicted to be restrictive and decisive for the function of the respective signaling (labeled in black). Other receptors (labeled in gray) are ignored at this time due to the following: 1. their lower abundance—only the top 150 highly expressed receptors from each cell type are considered; 2. less binding affinity—i.e. IGF1R vs INSR; 3. broad expression—i.e. LRP5 and FZD proteins are ubiquitously expressed across all cell types studied here; 4. known gene spatial expression—i.e. Fgfr2 is specifically expressed in alveolar epithelium. See Figure 4-Source Data 1 for the full list of these signaling genes selected for each cell type and from each dataset.

**Figure 4-Figure Supplement 2:**
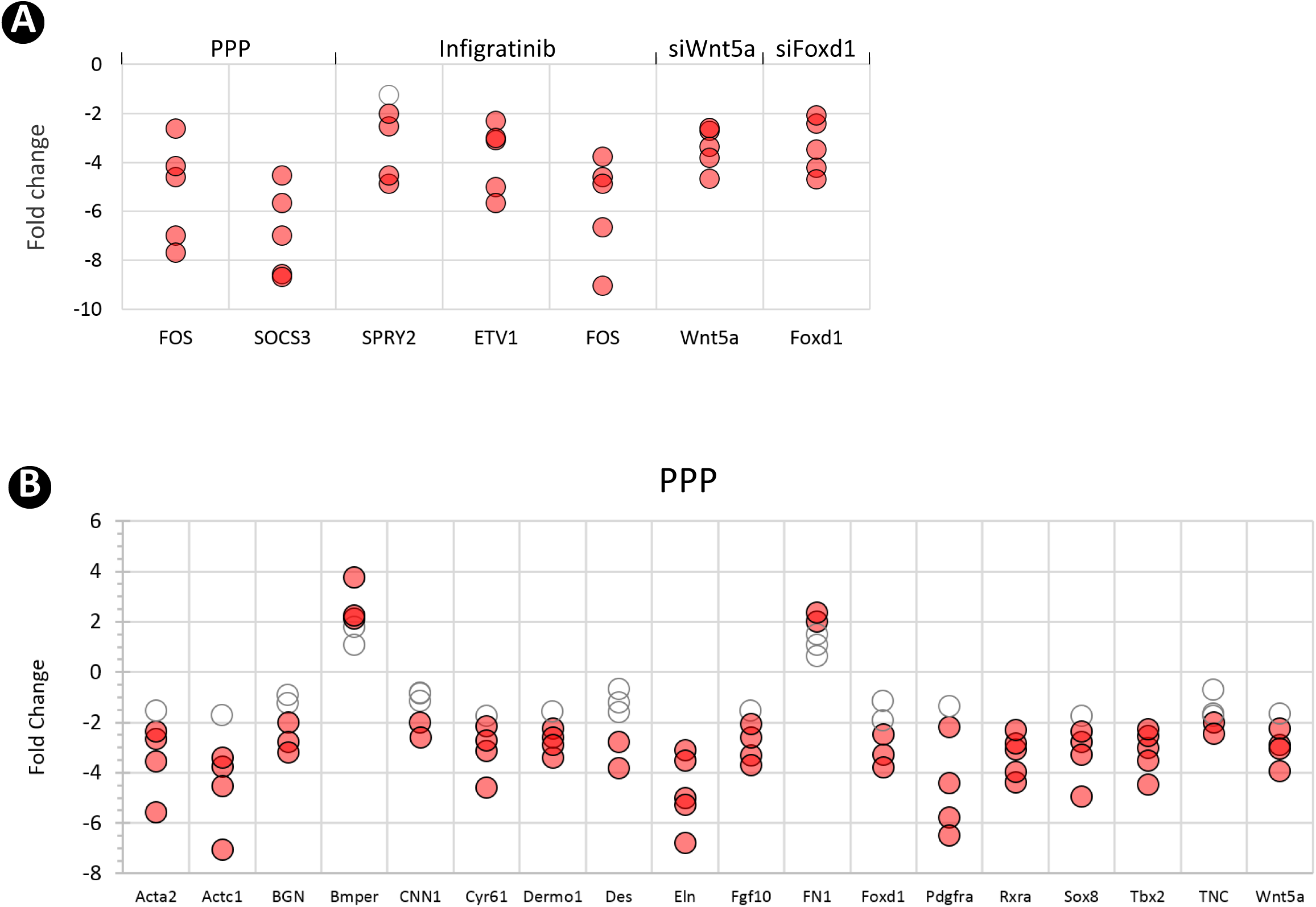
SCMF cell culture treatments. A: RT-PCR data showing the effect of various treatments (top) on their respective targeted genes (bottom). B: RT-PCR data from the altered SCMF genes demonstrating their response to the treatment from PPP.

Figure 4-Source Data 1: List of the ligands and receptors used for Secretome-Receptome analysis.

Figure 4-Source Data 2: List of the inhibitors and their concentrations used in cell culture treatment.

**Figure 5-Figure Supplement 1:**
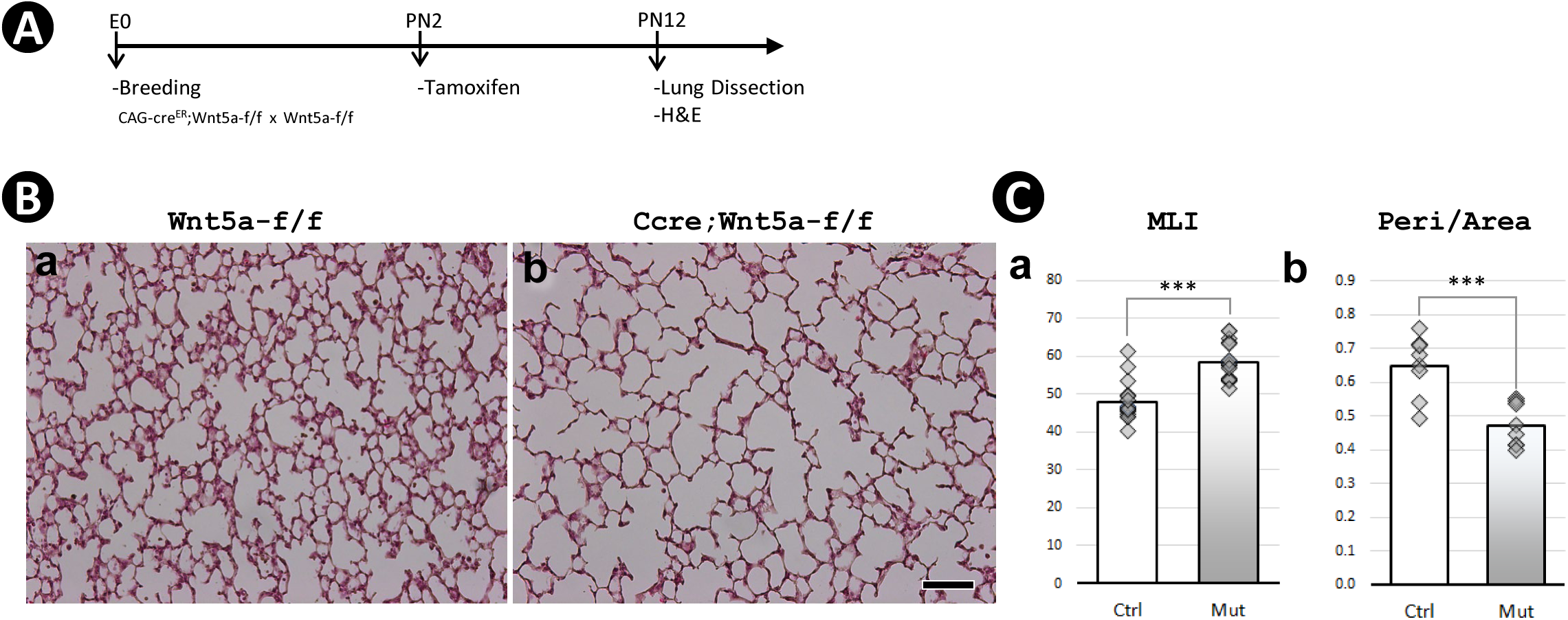
Postnatal inactivation of *Wnt5a*. A: Schematic of the experimental protocol. B: H&E staining of lung sections obtained from control (a) and Wnt5a-/- (b) mice as generated and described in Li *et al.,* 2020 and analyzed for this study. C: Morphometric measurements by MLI (c) and Peri/Area ratio (d). Scale bar: 100um for all images.

Figure 6-Source Data 1: List of clinical data for human neonatal lung samples.

## REFERENCES

1. Adam, M., Potter, A.S., Potter, S.S., 2017. Psychrophilic proteases dramatically reduce single-cell RNA-seq artifacts: a molecular atlas of kidney development. Development 144, 3625–3632.

2. Azeloglu, E.U., Iyengar, R., 2015. Signaling networks: information flow, computation, and decision making. Cold Spring Harb Perspect Biol 7, a005934.

3. Barber, R.D., Harmer, D.W., Coleman, R.A., Clark, B.J., 2005. GAPDH as a housekeeping gene: analysis of GAPDH mRNA expression in a panel of 72 human tissues. Physiol Genomics 21, 389–395.

4. Bostrom, H., Willetts, K., Pekny, M., Leveen, P., Lindahl, P., Hedstrand, H., Pekna, M., Hellstrom, M., Gebre-Medhin, S., Schalling, M., Nilsson, M., Kurland, S., Tornell, J., Heath, J.K., Betsholtz, C., 1996. PDGF-A signaling is a critical event in lung alveolar myofibroblast development and alveogenesis. Cell 85, 863–873.

5. Carre, C., Mas, A., Krouk, G., 2017. Reverse engineering highlights potential principles of large gene regulatory network design and learning. NPJ Syst Biol Appl 3, 17.

6. Chao, C.M., Moiseenko, A., Zimmer, K.P., Bellusci, S., 2016. Alveologenesis: key cellular players and fibroblast growth factor 10 signaling. Mol Cell Pediatr 3, 17.

7. Chao, C.M., Yahya, F., Moiseenko, A., Tiozzo, C., Shrestha, A., Ahmadvand, N., El Agha, E., Quantius, J., Dilai, S., Kheirollahi, V., Jones, M., Wilhem, J., Carraro, G., Ehrhardt, H., Zimmer, K.P., Barreto, G., Ahlbrecht, K., Morty, R.E., Herold, S., Abellar, R.G., Seeger, W., Schermuly, R., Zhang, J.S., Minoo, P., Bellusci, S., 2017. Fgf10 deficiency is causative for lethality in a mouse model of bronchopulmonary dysplasia. J Pathol 241, 91–103.

8. Chen, L., Chen, Q., Rong, P., Wang, H.Y., Chen, S., 2017. The energy sensing LKB1-AMPKalpha1 pathway regulates IGF1 secretion and consequent activation of the IGF1R-PKB pathway in primary hepatocytes. FEBS J 284, 2096–2109.

9. Cusanovich, D.A., Hill, A.J., Aghamirzaie, D., Daza, R.M., Pliner, H.A., Berletch, J.B., Filippova, G.N., Huang, X., Christiansen, L., DeWitt, W.S., Lee, C., Regalado, S.G., Read, D.F., Steemers, F.J., Disteche, C.M., Trapnell, C., Shendure, J., 2018. A Single-Cell Atlas of In Vivo Mammalian Chromatin Accessibility. Cell 174, 1309–1324 e1318.

10. Davidson, E.H., 2010. Emerging properties of animal gene regulatory networks. Nature 468, 911–920.

11. Davidson, E.H., Rast, J.P., Oliveri, P., Ransick, A., Calestani, C., Yuh, C.H., Minokawa, T., Amore, G., Hinman, V., Arenas-Mena, C., Otim, O., Brown, C.T., Livi, C.B., Lee, P.Y., Revilla, R., Rust, A.G., Pan, Z., Schilstra, M.J., Clarke, P.J., Arnone, M.I., Rowen, L., Cameron, R.A., McClay, D.R., Hood, L., Bolouri, H., 2002. A genomic regulatory network for development. Science 295, 1669–1678.

12. Dequeant, M.L., Pourquie, O., 2008. Segmental patterning of the vertebrate embryonic axis. Nat Rev Genet 9, 370–382.

13. Eng, C.L., Lawson, M., Zhu, Q., Dries, R., Koulena, N., Takei, Y., Yun, J., Cronin, C., Karp, C., Yuan, G.C., Cai, L., 2019. Transcriptome-scale super-resolved imaging in tissues by RNA seqFISH. Nature 568, 235–239.

14. Epaud, R., Aubey, F., Xu, J., Chaker, Z., Clemessy, M., Dautin, A., Ahamed, K., Bonora, M., Hoyeau, N., Flejou, J.F., Mailleux, A., Clement, A., Henrion-Caude, A., Holzenberger, M., 2012. Knockout of insulin-like growth factor-1 receptor impairs distal lung morphogenesis. PloS one 7, e48071.

15. Erwin, D.H., Davidson, E.H., 2009. The evolution of hierarchical gene regulatory networks. Nat Rev Genet 10, 141–148.

16. Gao, F., Davidson, E.H., 2008. Transfer of a large gene regulatory apparatus to a new developmental address in echinoid evolution. Proceedings of the National Academy of Sciences of the United States of America 105, 6091–6096.

17. Gao, F., Li, C., Danopoulos, S., Al Alam, D., Peinado, N., Webster, S., Borok, Z., Kohbodi, G.A., Bellusci, S., Minoo, P., 2022. Hedgehog-responsive PDGFRa(+) fibroblasts maintain a unique pool of alveolar epithelial progenitor cells during alveologenesis. Cell Rep 39, 110608.

18. Ghosh, M.C., Gorantla, V., Makena, P.S., Luellen, C., Sinclair, S.E., Schwingshackl, A., Waters, C.M., 2013. Insulin-like growth factor-I stimulates differentiation of ATII cells to ATI-like cells through activation of Wnt5a. Am J Physiol Lung Cell Mol Physiol 305, L222–228.

19. Hagan, A.S., Zhang, B., Ornitz, D.M., 2020. Identification of a FGF18-expressing alveolar myofibroblast that is developmentally cleared during alveologenesis. Development 147.

20. He, H., Snowball, J., Sun, F., Na, C.L., Whitsett, J.A., 2021. IGF1R controls mechanosignaling in myofibroblasts required for pulmonary alveologenesis. JCI Insight 6.

21. Jain, D., Bancalari, E., 2014. Bronchopulmonary dysplasia: clinical perspective. Birth Defects Res A Clin Mol Teratol 100, 134–144.

22. Jia, B., Xu, S., Xiao, G., Lamba, V., Liang, F., 2017. Learning gene regulatory networks from next generation sequencing data. Biometrics 73, 1221–1230.

23. Jin, K., Ruan, L., Pu, J., Zhong, A., Wang, F., Tan, S., Huang, H., Mu, J., Yang, G., 2018. Metformin suppresses growth and adrenocorticotrophic hormone secretion in mouse pituitary corticotroph tumor AtT20cells. Mol Cell Endocrinol 478, 53–61.

24. Jobe, A.J., 1999. The new BPD: an arrest of lung development. Pediatr Res 46, 641–643.

25. Juul, N.H., Stockman, C.A., Desai, T.J., 2020. Niche Cells and Signals that Regulate Lung Alveolar Stem Cells In Vivo. Cold Spring Harb Perspect Biol 12.

26. Kugler, M.C., Loomis, C.A., Zhao, Z., Cushman, J.C., Liu, L., Munger, J.S., 2017. Sonic Hedgehog Signaling Regulates Myofibroblast Function during Alveolar Septum Formation in Murine Postnatal Lung. Am J Respir Cell Mol Biol 57, 280–293.

27. Lau, L.F., 2011. CCN1/CYR61: the very model of a modern matricellular protein. Cell Mol Life Sci 68, 3149–3163.

28. Levine, M., Davidson, E.H., 2005. Gene regulatory networks for development. Proc Natl Acad Sci U S A 102, 4936-4942.

29. Ley, D., Hallberg, B., Hansen-Pupp, I., Dani, C., Ramenghi, L.A., Marlow, N., Beardsall, K., Bhatti, F., Dunger, D., Higginson, J.D., Mahaveer, A., Mezu-Ndubuisi, O.J., Reynolds, P., Giannantonio, C., van Weissenbruch, M., Barton, N., Tocoian, A., Hamdani, M., Jochim, E., Mangili, A., Chung, J.K., Turner, M.A., Smith, L.E.H., Hellstrom, A., study, t., 2019. rhIGF-1/rhIGFBP-3 in Preterm Infants: A Phase 2 Randomized Controlled Trial. J Pediatr 206, 56–65 e58.

30. Li, C., Lee, M.K., Gao, F., Webster, S., Di, H., Duan, J., Yang, C.Y., Bhopal, N., Peinado, N., Pryhuber, G., Smith, S.M., Borok, Z., Bellusci, S., Minoo, P., 2019. Secondary crest myofibroblast PDGFRalpha controls the elastogenesis pathway via a secondary tier of signaling networks during alveologenesis. Development 146.

31. Li, C., Li, M., Li, S., Xing, Y., Yang, C.Y., Li, A., Borok, Z., De Langhe, S., Minoo, P., 2015. Progenitors of secondary crest myofibroblasts are developmentally committed in early lung mesoderm. Stem Cells 33, 999–1012.

32. Li, C., Smith, S.M., Peinado, N., Gao, F., Li, W., Lee, M.K., Zhou, B., Bellusci, S., Pryhuber, G.S., Ho, H.H., Borok, Z., Minoo, P., 2020. WNT5a-ROR Signaling Is Essential for Alveologenesis. Cells 9. 999–1012.

33. Li, J., Yan, T., Wu, X., Ke, X., Li, X., Zhu, Y., Yang, J., Li, Z., 2021. Aberrant overexpression of transcription factor Forkhead box D1 predicts poor prognosis and promotes cancer progression in HNSCC. BMC Cancer 21, 1205.

34. Li, M., Li, C., Liu, Y.H., Xing, Y., Hu, L., Borok, Z., Kwong, K.Y., Minoo, P., 2008. Mesodermal deletion of transforming growth factor-beta receptor II disrupts lung epithelial morphogenesis: cross-talk between TGF-beta and Sonic hedgehog pathways. J Biol Chem 283, 36257–36264.

35. Li, R., Bernau, K., Sandbo, N., Gu, J., Preissl, S., Sun, X., 2018. Pdgfra marks a cellular lineage with distinct contributions to myofibroblasts in lung maturation and injury response. Elife 7.

36. Lim, W.A., Lee, C.M., Tang, C., 2013. Design principles of regulatory networks: searching for the molecular algorithms of the cell. Mol Cell 49, 202–212.

37. Longabaugh, W.J., 2012. BioTapestry: a tool to visualize the dynamic properties of gene regulatory networks. Methods Mol Biol 786, 359–394.

38. Longabaugh, W.J.R., Zeng, W., Zhang, J.A., Hosokawa, H., Jansen, C.S., Li, L., Romero-Wolf, M., Liu, P., Kueh, H.Y., Mortazavi, A., Rothenberg, E.V., 2017. Bcl11b and combinatorial resolution of cell fate in the T-cell gene regulatory network. Proc Natl Acad Sci U S A 114, 5800–5807.

39. Lopez, I.P., Pineiro-Hermida, S., Pais, R.S., Torrens, R., Hoeflich, A., Pichel, J.G., 2016. Involvement of Igf1r in Bronchiolar Epithelial Regeneration: Role during Repair Kinetics after Selective Club Cell Ablation. PloS one 11, e0166388.

40. Lopez, I.P., Rodriguez-de la Rosa, L., Pais, R.S., Pineiro-Hermida, S., Torrens, R., Contreras, J., Varela-Nieto, I., Pichel, J.G., 2015. Differential organ phenotypes after postnatal Igf1r gene conditional deletion induced by tamoxifen in UBC-CreERT2; Igf1r fl/fl double transgenic mice. Transgenic Res 24, 279–294.

41. Manchado, E., Weissmueller, S., Morris, J.P.t., Chen, C.C., Wullenkord, R., Lujambio, A., de Stanchina, E., Poirier, J.T., Gainor, J.F., Corcoran, R.B., Engelman, J.A., Rudin, C.M., Rosen, N., Lowe, S.W., 2016. A combinatorial strategy for treating KRAS-mutant lung cancer. Nature 534, 647–651.

42. Martin-Medina, A., Lehmann, M., Burgy, O., Hermann, S., Baarsma, H.A., Wagner, D.E., De Santis, M.M., Ciolek, F., Hofer, T.P., Frankenberger, M., Aichler, M., Lindner, M., Gesierich, W., Guenther, A., Walch, A., Coughlan, C., Wolters, P., Lee, J.S., Behr, J., Konigshoff, M., 2018. Increased Extracellular Vesicles Mediate WNT5A Signaling in Idiopathic Pulmonary Fibrosis. Am J Respir Crit Care Med 198, 1527–1538.

43. Materna, S.C., Oliveri, P., 2008. A protocol for unraveling gene regulatory networks. Nat Protoc 3, 1876–1887.

44. Mikels, A.J., Nusse, R., 2006. Purified Wnt5a protein activates or inhibits beta-catenin-TCF signaling depending on receptor context. PLoS Biol 4, e115.

45. Nabhan, A.N., Brownfield, D.G., Harbury, P.B., Krasnow, M.A., Desai, T.J., 2018. Single-cell Wnt signaling niches maintain stemness of alveolar type 2 cells. Science 359, 1118–1123.

46. Nakamura, H., Arai, Y., Totoki, Y., Shirota, T., Elzawahry, A., Kato, M., Hama, N., Hosoda, F., Urushidate, T., Ohashi, S., Hiraoka, N., Ojima, H., Shimada, K., Okusaka, T., Kosuge, T., Miyagawa, S., Shibata, T., 2015. Genomic spectra of biliary tract cancer. Nat Genet 47, 1003–1010.

47. Nakayama, S., Soejima, K., Yasuda, H., Yoda, S., Satomi, R., Ikemura, S., Terai, H., Sato, T., Yamaguchi, N., Hamamoto, J., Arai, D., Ishioka, K., Ohgino, K., Naoki, K., Betsuyaku, T., 2015. FOXD1 expression is associated with poor prognosis in non-small cell lung cancer. Anticancer Res 35, 261–268.

48. Narasaraju, T.A., Chen, H., Weng, T., Bhaskaran, M., Jin, N., Chen, J., Chen, Z., Chinoy, M.R., Liu, L., 2006. Expression profile of IGF system during lung injury and recovery in rats exposed to hyperoxia: a possible role of IGF-1 in alveolar epithelial cell proliferation and differentiation. J Cell Biochem 97, 984–998.

49. Negretti, N.M., Plosa, E.J., Benjamin, J.T., Schuler, B.A., Habermann, A.C., Jetter, C.S., Gulleman, P., Bunn, C., Hackett, A.N., Ransom, M., Taylor, C.J., Nichols, D., Matlock, B.K., Guttentag, S.H., Blackwell, T.S., Banovich, N.E., Kropski, J.A., Sucre, J.M.S., 2021. A single-cell atlas of mouse lung development. Development 148.

50. Nemoto, E., Ebe, Y., Kanaya, S., Tsuchiya, M., Nakamura, T., Tamura, M., Shimauchi, H., 2012. Wnt5a signaling is a substantial constituent in bone morphogenetic protein-2-mediated osteoblastogenesis. Biochem Biophys Res Commun 422, 627–632.

51. Olson, E.N., 2006. Gene regulatory networks in the evolution and development of the heart. Science 313, 1922–1927.

52. Peter, I.S., Davidson, E.H., 2009. Modularity and design principles in the sea urchin embryo gene regulatory network. FEBS Lett 583, 3948–3958.

53. Peter, I.S., Davidson, E.H., 2017. Assessing regulatory information in developmental gene regulatory networks. Proc Natl Acad Sci U S A 114, 5862–5869.

54. Peter, I.S., Faure, E., Davidson, E.H., 2012. Predictive computation of genomic logic processing functions in embryonic development. Proc Natl Acad Sci U S A 109, 16434–16442.

55. Pires-daSilva, A., Sommer, R.J., 2003. The evolution of signaling pathways in animal development. Nat Rev Genet 4, 39–49.

56. Przybyla, L., Gilbert, L.A., 2021. A new era in functional genomics screens. Nat Rev Genet.

57. Ramilowski, J.A., Goldberg, T., Harshbarger, J., Kloppmann, E., Lizio, M., Satagopam, V.P., Itoh, M., Kawaji, H., Carninci, P., Rost, B., Forrest, A.R., 2015. A draft network of ligand-receptor-mediated multicellular signaling in human. Nat Commun 6, 7866.

58. Royo, J.L., Maeso, I., Irimia, M., Gao, F., Peter, I.S., Lopes, C.S., D’Aniello, S., Casares, F., Davidson, E.H., Garcia-Fernandez, J., Gomez-Skarmeta, J.L., 2011. Transphyletic conservation of developmental regulatory state in animal evolution. Proceedings of the National Academy of Sciences of the United States of America 108, 14186–14191.

59. Sakisaka, Y., Tsuchiya, M., Nakamura, T., Tamura, M., Shimauchi, H., Nemoto, E., 2015. Wnt5a attenuates Wnt3a-induced alkaline phosphatase expression in dental follicle cells. Exp Cell Res 336, 85–93.

60. Sanson, K.R., Hanna, R.E., Hegde, M., Donovan, K.F., Strand, C., Sullender, M.E., Vaimberg, E.W., Goodale, A., Root, D.E., Piccioni, F., Doench, J.G., 2018. Optimized libraries for CRISPR-Cas9 genetic screens with multiple modalities. Nat Commun 9, 5416.

61. Satou, Y., Satoh, N., Imai, K.S., 2009. Gene regulatory networks in the early ascidian embryo. Biochim Biophys Acta 1789, 268–273.

62. Sauka-Spengler, T., Bronner-Fraser, M., 2008. A gene regulatory network orchestrates neural crest formation. Nat Rev Mol Cell Biol 9, 557–568.

63. Schuler, B.A., Habermann, A.C., Plosa, E.J., Taylor, C.J., Jetter, C., Negretti, N.M., Kapp, M.E., Benjamin, J.T., Gulleman, P., Nichols, D.S., Braunstein, L.Z., Hackett, A., Koval, M., Guttentag, S.H., Blackwell, T.S., Webber, S.A., Banovich, N.E., Vanderbilt, C.-C.C., Human Cell Atlas Biological, N., Kropski, J.A., Sucre, J.M., 2021. Age-determined expression of priming protease TMPRSS2 and localization of SARS-CoV-2 in lung epithelium. J Clin Invest 131.

64. Seedorf, G., Kim, C., Wallace, B., Mandell, E.W., Nowlin, T., Shepherd, D., Abman, S.H., 2020. rhIGF-1/BP3 Preserves Lung Growth and Prevents Pulmonary Hypertension in Experimental Bronchopulmonary Dysplasia. Am J Respir Crit Care Med 201, 1120–1134.

65. Short, E.J., Kirchner, H.L., Asaad, G.R., Fulton, S.E., Lewis, B.A., Klein, N., Eisengart, S., Baley, J., Kercsmar, C., Min, M.O., Singer, L.T., 2007. Developmental sequelae in preterm infants having a diagnosis of bronchopulmonary dysplasia: analysis using a severity-based classification system. Arch Pediatr Adolesc Med 161, 1082–1087.

66. Skene, P.J., Henikoff, S., 2015. A simple method for generating high-resolution maps of genome-wide protein binding. Elife 4, e09225.

67. Sucre, J.M.S., Vickers, K.C., Benjamin, J.T., Plosa, E.J., Jetter, C.S., Cutrone, A., Ransom, M., Anderson, Z., Sheng, Q., Fensterheim, B.A., Ambalavanan, N., Millis, B., Lee, E., Zijlstra, A., Konigshoff, M., Blackwell, T.S., Guttentag, S.H., 2020. Hyperoxia Injury in the Developing Lung Is Mediated by Mesenchymal Expression of Wnt5A. Am J Respir Crit Care Med 201, 1249–1262.

68. Sun, X., Perl, A.K., Li, R., Bell, S.M., Sajti, E., Kalinichenko, V.V., Kalin, T.V., Misra, R.S., Deshmukh, H., Clair, G., Kyle, J., Crotty Alexander, L.E., Masso-Silva, J.A., Kitzmiller, J.A., Wikenheiser-Brokamp, K.A., Deutsch, G., Guo, M., Du, Y., Morley, M.P., Valdez, M.J., Yu, H.V., Jin, K., Bardes, E.E., Zepp, J.A., Neithamer, T., Basil, M.C., Zacharias, W.J., Verheyden, J., Young, R., Bandyopadhyay, G., Lin, S., Ansong, C., Adkins, J., Salomonis, N., Aronow, B.J., Xu, Y., Pryhuber, G., Whitsett, J., Morrisey, E.E., Consortium, N.L., 2022. A census of the lung: CellCards from LungMAP. Dev Cell 57, 112–145 e112.

69. Tsao, P.N., Matsuoka, C., Wei, S.C., Sato, A., Sato, S., Hasegawa, K., Chen, H.K., Ling, T.Y., Mori, M., Cardoso, W.V., Morimoto, M., 2016. Epithelial Notch signaling regulates lung alveolar morphogenesis and airway epithelial integrity. Proc Natl Acad Sci U S A 113, 8242–8247.

70. Verheyden, J.M., Sun, X., 2020. A transitional stem cell state in the lung. Nat Cell Biol 22, 1025–1026.

71. Wang, G., Yin, L., Peng, Y., Gao, Y., Gao, H., Zhang, J., Lv, N., Miao, Y., Lu, Z., 2019. Insulin promotes invasion and migration of KRAS(G12D) mutant HPNE cells by upregulating MMP-2 gelatinolytic activity via ERK- and PI3K-dependent signalling. Cell Prolif 52, e12575.

72. Wang, Y., Tang, Z., Huang, H., Li, J., Wang, Z., Yu, Y., Zhang, C., Li, J., Dai, H., Wang, F., Cai, T., Tang, N., 2018a. Pulmonary alveolar type I cell population consists of two distinct subtypes that differ in cell fate. Proc Natl Acad Sci U S A 115, 2407–2412.

73. Wang, Z., Li, W., Guo, Q., Wang, Y., Ma, L., Zhang, X., 2018b. Insulin-Like Growth Factor-1 Signaling in Lung Development and Inflammatory Lung Diseases. Biomed Res Int 2018, 6057589.

74. Wong, G.S., Zhou, J., Liu, J.B., Wu, Z., Xu, X., Li, T., Xu, D., Schumacher, S.E., Puschhof, J., McFarland, J., Zou, C., Dulak, A., Henderson, L., Xu, P., O’Day, E., Rendak, R., Liao, W.L., Cecchi, F., Hembrough, T., Schwartz, S., Szeto, C., Rustgi, A.K., Wong, K.K., Diehl, J.A., Jensen, K., Graziano, F., Ruzzo, A., Fereshetian, S., Mertins, P., Carr, S.A., Beroukhim, R., Nakamura, K., Oki, E., Watanabe, M., Baba, H., Imamura, Y., Catenacci, D., Bass, A.J., 2018. Targeting wild-type KRAS-amplified gastroesophageal cancer through combined MEK and SHP2 inhibition. Nat Med 24, 968–977.

75. Wu, H., Larribere, L., Sun, Q., Novak, D., Sachindra, S., Granados, K., Umansky, V., Utikal, J., 2018. Loss of neural crest-associated gene FOXD1 impairs melanoma invasion and migration via RAC1B downregulation. Int J Cancer 143, 2962–2972.

76. Wu, H., Tang, N., 2021. Stem cells in pulmonary alveolar regeneration. Development 148.

77. Xu, Y., Wang, Y., Besnard, V., Ikegami, M., Wert, S.E., Heffner, C., Murray, S.A., Donahue, L.R., Whitsett, J.A., 2012. Transcriptional programs controlling perinatal lung maturation. PloS one 7, e37046.

78. Zepp, J.A., Morley, M.P., Loebel, C., Kremp, M.M., Chaudhry, F.N., Basil, M.C., Leach, J.P., Liberti, D.C., Niethamer, T.K., Ying, Y., Jayachandran, S., Babu, A., Zhou, S., Frank, D.B., Burdick, J.A., Morrisey, E.E., 2021. Genomic, epigenomic, and biophysical cues controlling the emergence of the lung alveolus. Science 371.

79. Zepp, J.A., Morrisey, E.E., 2019. Cellular crosstalk in the development and regeneration of the respiratory system. Nat Rev Mol Cell Biol 20, 551–566.

80. Zhang, K., Yao, E., Lin, C., Chou, Y.T., Wong, J., Li, J., Wolters, P.J., Chuang, P.T., 2020. A mammalian Wnt5a-Ror2-Vangl2 axis controls the cytoskeleton and confers cellular properties required for alveologenesis. Elife 9.

81. Zhao, G., Cheng, X.W., Piao, L., Hu, L., Lei, Y., Yang, G., Inoue, A., Ogasawara, S., Wu, H., Hao, C.N., Okumura, K., Kuzuya, M., 2017. The Soluble VEGF Receptor sFlt-1 Contributes to Impaired Neovascularization in Aged Mice. Aging Dis 8, 287–300.

